# The Bromodomains of the mammalian SWI/SNF (mSWI/SNF) ATPases Brahma (BRM) and Brahma Related Gene 1 (BRG1) promote chromatin interaction and are critical for skeletal muscle differentiation

**DOI:** 10.1101/2020.08.25.267666

**Authors:** Tapan Sharma, Hanna Witwicka, Anthony N. Imbalzano

## Abstract

Skeletal muscle differentiation induces changes in the epigenome of myoblasts as they proceed towards a myogenic phenotype. mSWI/SNF chromatin remodeling enzymes coordinate with lineage-determining transcription factors and are key regulators of differentiation. Three mSWI/SNF proteins, the mutually exclusive ATPases, BRG1 and BRM, and the BAF180 protein (Polybromo1, PBRM1) contain bromodomains belonging to the same structural subfamily. Bromodomains bind to acetylated lysines on histone N-terminal tails and on other proteins. Pharmacological inhibition of mSWI/SNF bromodomain function using the selective inhibitor PFI-3 reduced differentiation, decreased expression of myogenic genes, and increased the expression of cell cycle-related genes, and the number of cells that remained in the cell cycle. Knockdown of BAF180 had no effect on differentiation, suggesting that only the BRG1 and BRM bromodomains contributed to differentiation. Comparison with existing gene expression data from myoblasts subjected to knockdown of BRG1 or BRM showed that bromodomain function was required for a subset of BRG1- and BRM-dependent gene expression. ChIP analysis revealed decreased BRG1 and BRM binding to target gene promoters, indicating that the BRG1 and BRM bromodomains promote chromatin binding. Thus mSWI/SNF ATPase bromodomains contribute to cell cycle exit, to skeletal muscle-specific gene expression, and to stable promoter binding by the mSWI/SNF ATPases.

## INTRODUCTION

Regulation of gene expression is a tightly coordinated process that is dependent on transcription factors, coactivators and chromatin remodelers. Some of these regulators are tissue-specific and act on target genes in a context-dependent manner. Tissue-specific regulation is absolutely crucial for proper development of multi-cellular life forms in which all cells contain the same genetic information. Portions of the genome that are irrelevant to a particular tissue type are often condensed into repressive heterochromatin as development and differentiation occur (1, 2). In contrast, coordinated activity of lineage-determining transcription factors and chromatin remodelers, in particular the mSWI/SNF family of chromatin remodeling enzymes, drives many differentiation events, including skeletal muscle differentiation (3–7). The mSWI/SNF enzymes remodel chromatin in an ATP-dependent manner (8–10) and form a family of enzymes assembled into different configurations from a potential pool of more than twenty subunit proteins (11–13). The BRG1 and BRM ATPases act as mutually exclusive catalytic subunits (10).

Skeletal muscle originates from the paraxial mesoderm during embryogenesis. Fetal skeletal myogenesis is characterized by an abundance of myogenic progenitor cells that divide actively and fuse to form multinucleated muscle fibers (14, 15). As the embryo develops into an adult, these progenitor cells become relatively sparse and quiescent. These adult stem cells are known as satellite cells and can be activated to proliferate and regenerate new myofibers in case of an injury to adult skeletal muscle (16–18). Upon activation, expression of myogenic regulatory factors (MRFs) – MYOD, MRF4, MYF5 and Myogenin - is initiated in a coordinated manner. MRFs are basic helix-loop-helix (bHLH) proteins that are evolutionarily conserved from worms to humans (19–21). They bind to consensus sequences called E-boxes at target muscle promoters and activate muscle-specific gene expression (22). Another family of transcription factors called the myocyte enhancer factor 2 (MEF2) family acts with the MRFs to promote expression of the myogenic genes (23, 24).

During skeletal myogenesis, the mSWI/SNF complex is recruited to the myogenic loci by MRFs (25–30), in some cases, in conjunction with PBX1 (25). Mechanistically, upon induction of differentiation in myocytes, the p38 kinase responds to extracellular cues by phosphorylating the BAF60c subunit of mSWI/SNF chromatin remodeling enzymes, which is associated with MYOD on myogenic genes in the absence of other mSWI/SNF subunits in proliferating myoblasts (27, 29). The phospho-BAF60c-MYOD complex then recruits the rest of the mSWI/SNF complex to myogenic loci, which promotes chromatin accessibility and activates gene expression (29). Once recruited to myogenic loci, the ATPase activity of BRG1 or BRM in the complex is known to be indispensable for expression of the differentiation-specific gene program (26, 28, 31).

The BRG1 and BRM ATPases possess bromodomains in the C-terminal part of the protein (32–34). Bromodomains are well-characterized motifs known to interact with acetylated lysine residues on the N-terminal tails of histones H3 and H4 (35, 36) and on other non-histone proteins (37). The interaction of bromodomains with acetylated histones has been determined to be crucial for regulation of some gene expression events (37). Based on structural homology, bromodomain-containing proteins can be classified into eight families (35, 38). BRG1 and BRM belong to family VIII of bromodomains along with a third mSWI/SNF protein called BAF180 (Polybromo1, PBRM1, PB1) that contains six tandem bromodomains (38, 39)

In this study, we characterized the role of mSWI/SNF bromodomains in the context of skeletal myogenesis. We showed that inhibiting bromodomain function using PFI-3, a specific pharmacological inhibitor that binds to the BRG1, BRM and BAF180 bromodomains (40–42), reduced the ability of mouse myoblasts to differentiate into myotubes. Using RNA-sequencing, we identified the genes whose expression is dependent on mSWI/SNF bromodomains. Broadly, proliferation-related genes were found to be upregulated by bromodomain inhibition while myogenic genes were downregulated. We also demonstrated that bromodomain function is essential for timely exit of myoblasts from the cell cycle upon induction of differentiation. We determined that BAF180 is not required for myogenesis in mouse myoblasts and demonstrated that the BRG1 and BRM bromodomains play a crucial role in skeletal muscle differentiation by promoting the stable binding of BRG1 and BRM to target gene promoters. Thus, this study mechanistically demonstrates the specific importance of mSWI/SNF bromodomains in context of skeletal muscle differentiation.

## RESULTS

### Inhibition of bromodomain function results in aberrant myotube fusion

PFI-3 is a pharmacological inhibitor specific for the BRG1, BRM and BAF180 bromodomains, members of bromodomain family VIII (40, 41). Prior work showed that PFI-3 impaired differentiation of immortalized pre-adipocytes and myoblasts (42). The mechanisms responsible for the observed effects on differentiation were not defined, so we sought to investigate the roles played by mSWI/SNF bromodomains during myogenesis.

C2C12 immortalized myoblasts and primary myoblasts isolated from the tibialis anterior muscles of 1-week old C57BL/6 mice were assayed for their ability to differentiate in the presence of PFI-3 or the vehicle (DMSO). DMSO-treated C2C12 myoblasts immunostained for myosin heavy chain (MHC) showed formation of longer and thicker myotubes at 48h and 72h post induction of differentiation than did C2C12 myoblasts treated with PFI-3 (Fig 1a). The efficiency of myogenic differentiation can be scored by calculating fusion index, which is the ratio of the number of nuclei in MHC-stained cells to the total number of nuclei (43). C2C12 cells treated with PFI-3 showed a >50% decrease in fusion index at 24h, 48h and 72h as compared to control samples (Fig 1b). Similar results were observed when primary myoblasts were exposed to PFI-3. While DMSO-treated primary myoblasts showed elongated myotubes upon induction of differentiation, the PFI-3-treated cells showed fewer and less elongated myotubes at corresponding timepoints (Supp fig 1a). Quantitative analysis of differentiated primary myoblasts immunostained for MHC showed about a 25-30% decrease in fusion index (Supp fig 1b).

**Fig. 1.**
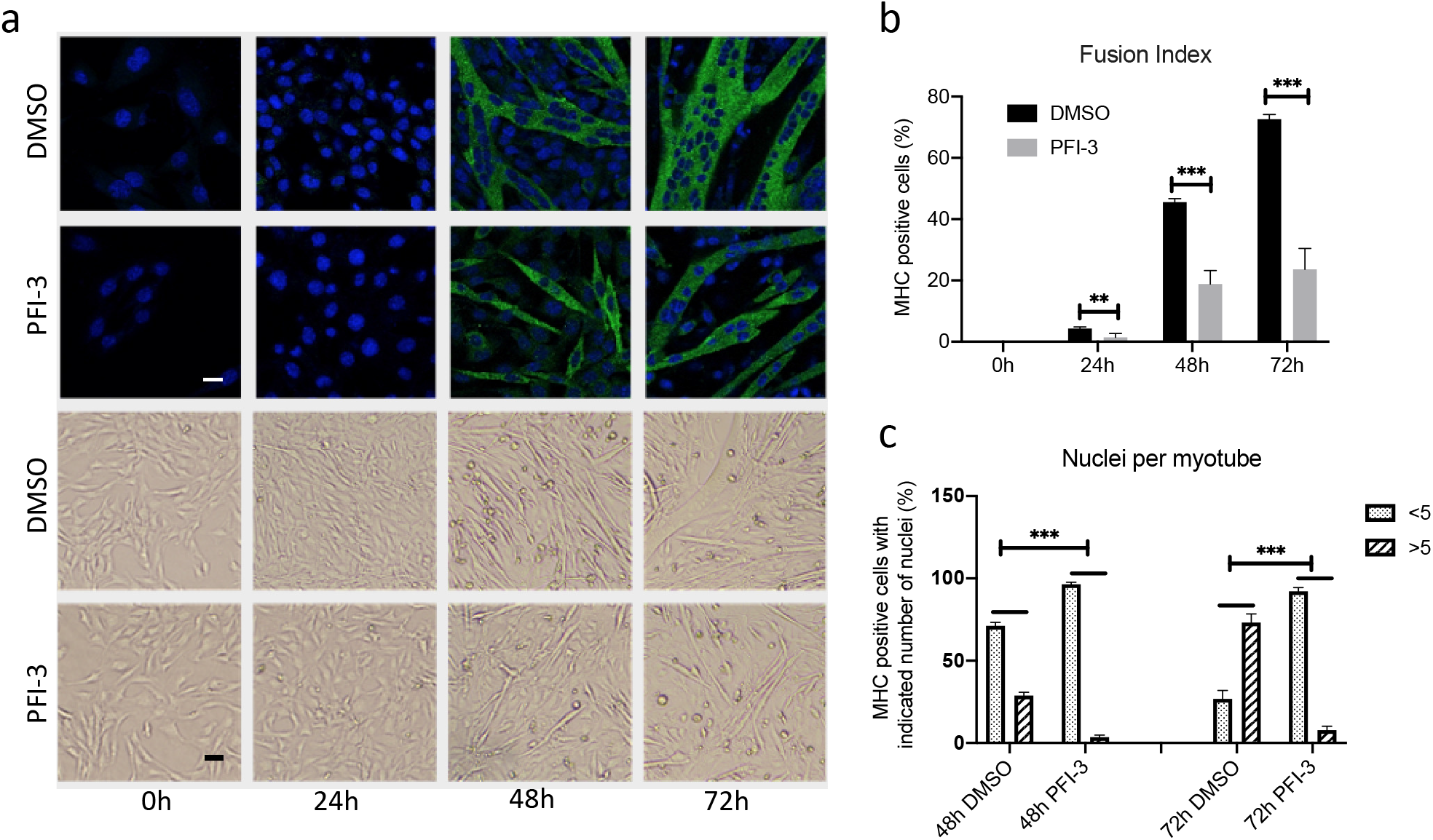
(a) Confocal (top, scale 5μm) and bright field (bottom, scale 20μm) images for C2C12 myoblasts treated with DMSO or PFI-3 and stained for myosin heavy chain (green) and with DAPI (blue) at the indicated timepoints. (b) Quantification of fusion index. (c) Differentiated myoblasts at respective timepoints were analyzed for number of nuclei per myotube. *p<0.05, **p<0.01 and ***p<0.005

We further quantitatively analyzed the extent of differentiation by counting the number of nuclei in MHC-positive myotubes and classifying them into groups at each timepoint. DMSO-treated C2C12 cells shifted from the majority of 48h myotubes having <5 nuclei to the majority of myotubes having >5 nuclei by 72h (Fig 1c). PFI-3-treated cells failed to make this switch; the majority of 72h myotubes had <5 nuclei (Fig 1c). Similarly, in PFI-3 treated primary cells, the number of myotubes with >5 nuclei at 36h was about one-third of those in DMSO-treated control cells. In PFI-3 treated samples, cells with a single nucleus positively immunostained for MHC were abundant, showing a failure of differentiating myoblasts to fuse (Supp fig 1c). These results suggest that an initial myogenic stimulus is present but is not fully implemented due to inhibition of bromodomain function.

### Myogenic genes are downregulated upon PFI-3 induced inhibition of mSWI/SNF bromodomains

The results show that PFI-3 treatment causes defects in myogenic differentiation, including an inability of the differentiating myoblasts and/or nascent myotubes to fuse. Myomaker and myomixer have been identified as master regulators of myoblast fusion (44–47). We therefore determined whether the expression of these two regulators was altered upon PFI-3 induced bromodomain inhibition. The results show that expression of these two genes was significantly lower in PFI-3 treated C2C12 cells (Fig 2a). The expression of other myogenic genes like myogenin, creatine kinase and myosin light chain 1 was also significantly decreased in PFI-3 treated samples, as was the expression of caveolin 3 and integrin 7A, two muscle differentiation-related genes (Fig 2b). Western blot analysis confirmed the decreased expression of myosin heavy chain in PFI-3 treated C2C12 cells (Fig 2c). Similar results were obtained for PFI-3 treated primary myoblasts (Supp fig 2). The gene expression signatures from both C2C12 cells and primary myoblasts provides a molecular explanation for the differentiation phenotype caused by bromodomain inhibition.

**Fig. 2.**
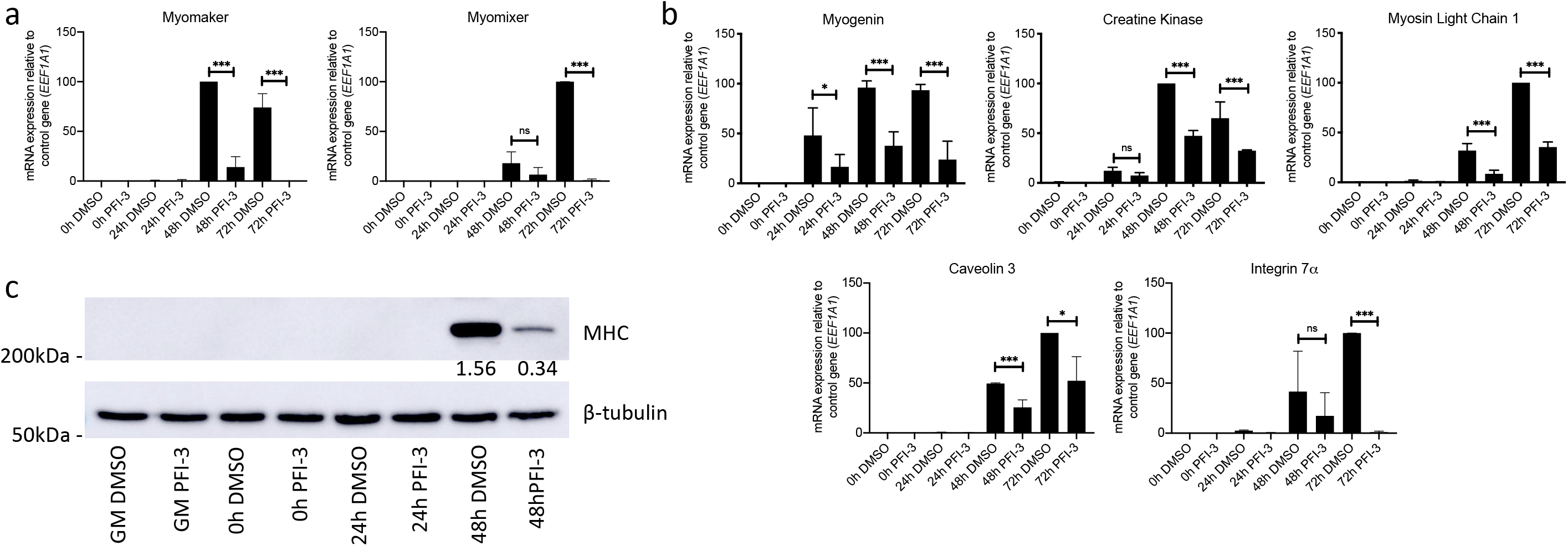
mRNA expression levels at the indicated timepoints for (a) the fusion regulator genes myomaker and myomixer and (b) the myogenic genes myogenin, muscle creatine kinase, myosin light chain 1, Caveolin 3 and Integrin 7α in C2C12 myoblasts treated with DMSO or PFI-3. Expression was normalized to a control gene (EEF1A1). 100% expression is defined as the timepoint at which maximal expression was observed. ns, not significant, *p<0.05, **p<0.01 and ***p<0.005. (c) Representative western blot for MHC expression at the indicated times in C2C12 cells treated with DMSO or PFI-3. The indicated numbers are the pixel counts normalized to β-tubulin expression calculated using ImageJ.

### RNA-seq analysis of PFI-3 treated C2C12 cells shows upregulation of cell cycle genes and downregulation of myogenic genes

To gain better insight into effect of the molecular mechanism of bromodomain inhibition on skeletal muscle differentiation, we performed RNA-sequencing of C2C12 cells treated with DMSO or PFI-3. Cells were harvested from proliferative stage (GM) and two differentiated stages (DM 24h and DM 48h post-induction). Libraries generated from the samples had ~45M unique reads. Transcripts were mapped to the mouse genome (mm10) and gene expression levels were calculated. Genes that were identified to be differentially expressed in both replicates for each condition and timepoint were considered for further analysis.

We first examined whether PFI-3 treatment affected gene expression of the subunits of mSWI/SNF complexes. A recent characterization of sub-families of mSWI/SNF complexes identified 29 subunit proteins (13). Assessment of expression of the genes encoding each of these proteins at each time point found only two instances of statistically significant differences (Supp. Table 1). *Arid1a* expression was reduced ~7% at 24 h post-differentiation and *Actl6a* expression was increased ~27% at 48 h post-differentiation. We conclude that PFI-3 treatment had essentially no effect on the expression of the genes encoding mSWI/SNF subunits.

Inhibition of bromodomain function affected the expression of about 50% of the total genes identified as expressed over the time course of the experiment (Fig 3a). The number of DEGs due to bromodomain inhibition increased as a function of differentiation (Fig 3b). The total number of DEGs for proliferating cells (GM) and differentiating cells at 24h or 48h post-differentiation (DM 24h and DM 48h) were 3144 (up 2216; down 928), 4675 (up 2878; down 1797) and 5261 (up 3105; down 2156), respectively (Supp. Table 2). Gene expression at DM 24h and DM 48h was strongly correlated with 2359 common DEGs as compared to about 1634 common DEGs between GM and DM 24h. There were 899 genes that were differentially expressed at all timepoints. Gene ontology (GO) analysis was performed on DEGs to cluster genes into function-based categories (48, 49) and the complete results are listed in Supp. Table 3. GO analysis of genes downregulated at DM 48h showed that the top 10 biological process categories were related to skeletal muscle contraction and skeletal muscle tissue development (Fig 3c; Supp. Table 3). This is in agreement with our experimental results, thus identifying the importance of bromodomain function in myogenesis. The top 10 categories from GO analysis of genes upregulated at DM 48h were related to cell proliferation (Fig 3d; Supp. Table 3), which indicated altered proliferation due to PFI-3-induced bromodomain inhibition. The promoters of the differentially expressed genes were also analyzed using the HOMER motif enrichment software (49). Sequences 1kb upstream of the transcription start sites were searched for presence of known consensus motifs (Supp. Table 4). The analysis revealed that promoters of genes downregulated due to PFI-3 treatment were significantly enriched in motifs corresponding to muscle specific transcription factors from the MEF and MRF families. (Fig 3c; Supp. Table 4). In the case of upregulated genes, HOMER analysis identified enrichment of motifs known to be bound by E2F family, NFY, KLF5 and Sp1 transcription factors (Fig 3d; Supp. Table 4). E2F and KLF5 TF families are known to play key role in regulation of cell proliferation and differentiation (50–52). Thus, PFI-3 induced bromodomain inhibition affects expression of genes which are involved in regulation of cell proliferation and skeletal muscle differentiation.

**Fig. 3.**
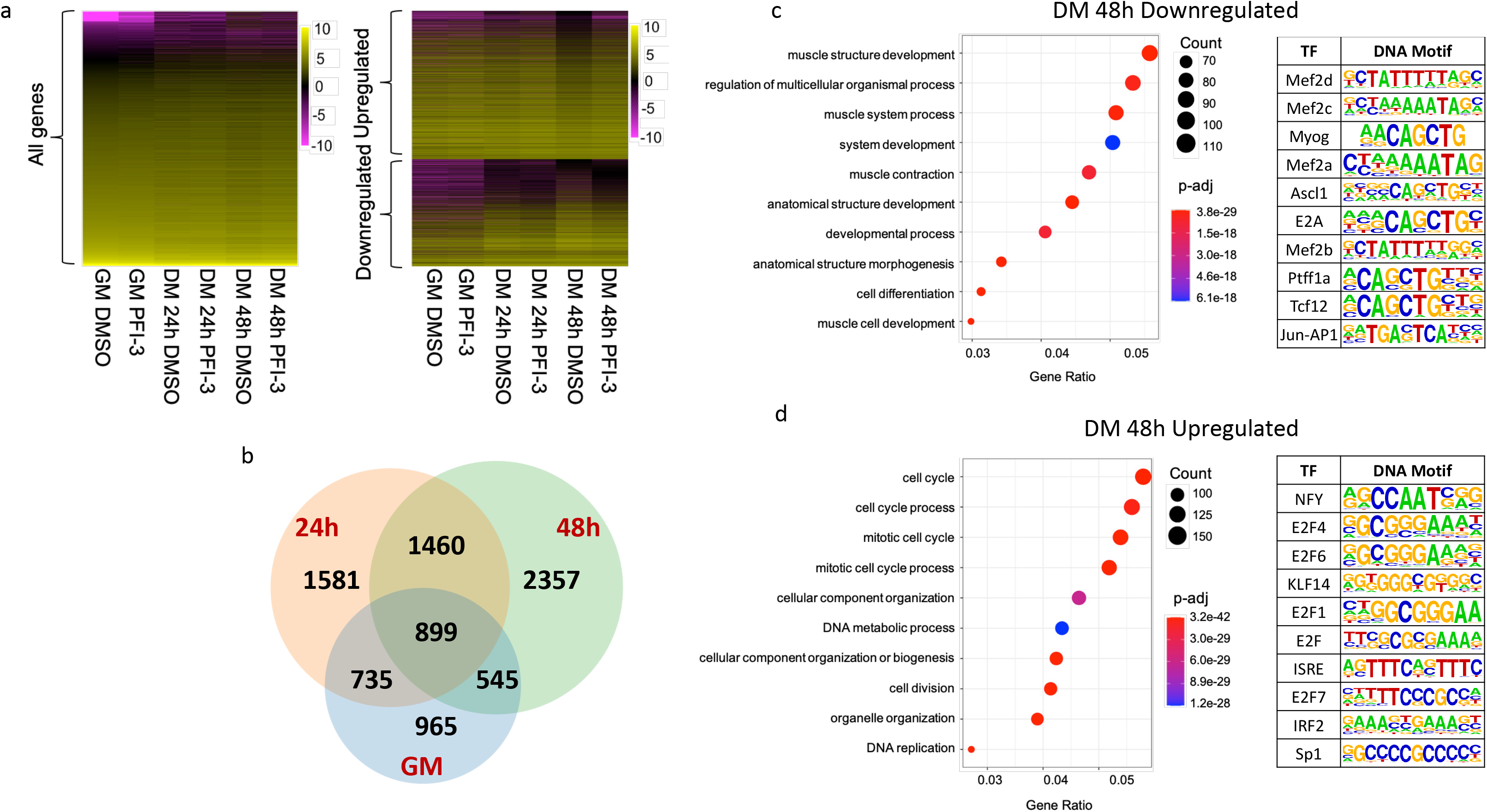
(a) Heat maps showing results from RNA-seq analysis of PFI-3 treated C2C12 cells assayed while in the proliferative stage in growth media (GM) and while in differentiation media (DM) for 24h and 48h. (b) Venn diagram showing the number of genes affected at different timepoints. There were 572 differentially expressed genes (DEGs) in GM (blue), 1319 DEGs in DM 24h (peach) and 1681 DEGs in DM 48h (green). (c) GO analysis of downregulated genes at 48h post induction of differentiation shows downregulation of muscle related genes. A HOMER motif search shows enrichment of motifs corresponding to muscle-specific transcription factor families. (d) GO analysis of upregulated genes at 48h post induction of differentiation shows upregulation of cell cycle related processes. A HOMER motif search shows enrichment of transcription factor motifs associated with cell cycle regulators.

### PFI-3 treatment blocked cell-cycle exit of C2C12 cells induced for differentiation

Cell cycle exit is prerequisite for cellular differentiation to proceed in a number of cell types (53–55). Results from the GO analysis of upregulated genes at DM 48h indicated that inhibition of bromodomain function may interfere with cell cycle exit. To experimentally address the requirement of mSWI/SNF bromodomain function in cell cycle exit, a BrdU (5-bromo-deoxyuridine) incorporation assay was performed with samples treated with or without PFI-3 (Fig 4a). Confocal microscopy analysis showed that cells treated with PFI-3 continued to incorporate BrdU even after the control cells showed no further incorporation, indicating a partial inability to exit cell cycle. Quantification of these images is shown in Fig 4b. Increased mRNA expression of cyclin A2, cyclin B1, cyclin D1, and cyclin D2 in PFI-3 treated samples from DM 48h as compared to DMSO controls further correlates with continued cell cycle (Fig 4c). These results show that bromodomain inhibition allows some of the myoblasts to overcome the signals to exit cycle that are normally provided by the low mitogen media and by contact inhibition. Thus, PFI-3 induced bromodomain inhibition may be affecting two aspects of myogenesis: timely exit from the cell cycle and the expression of myogenic genes.

**Fig. 4.**
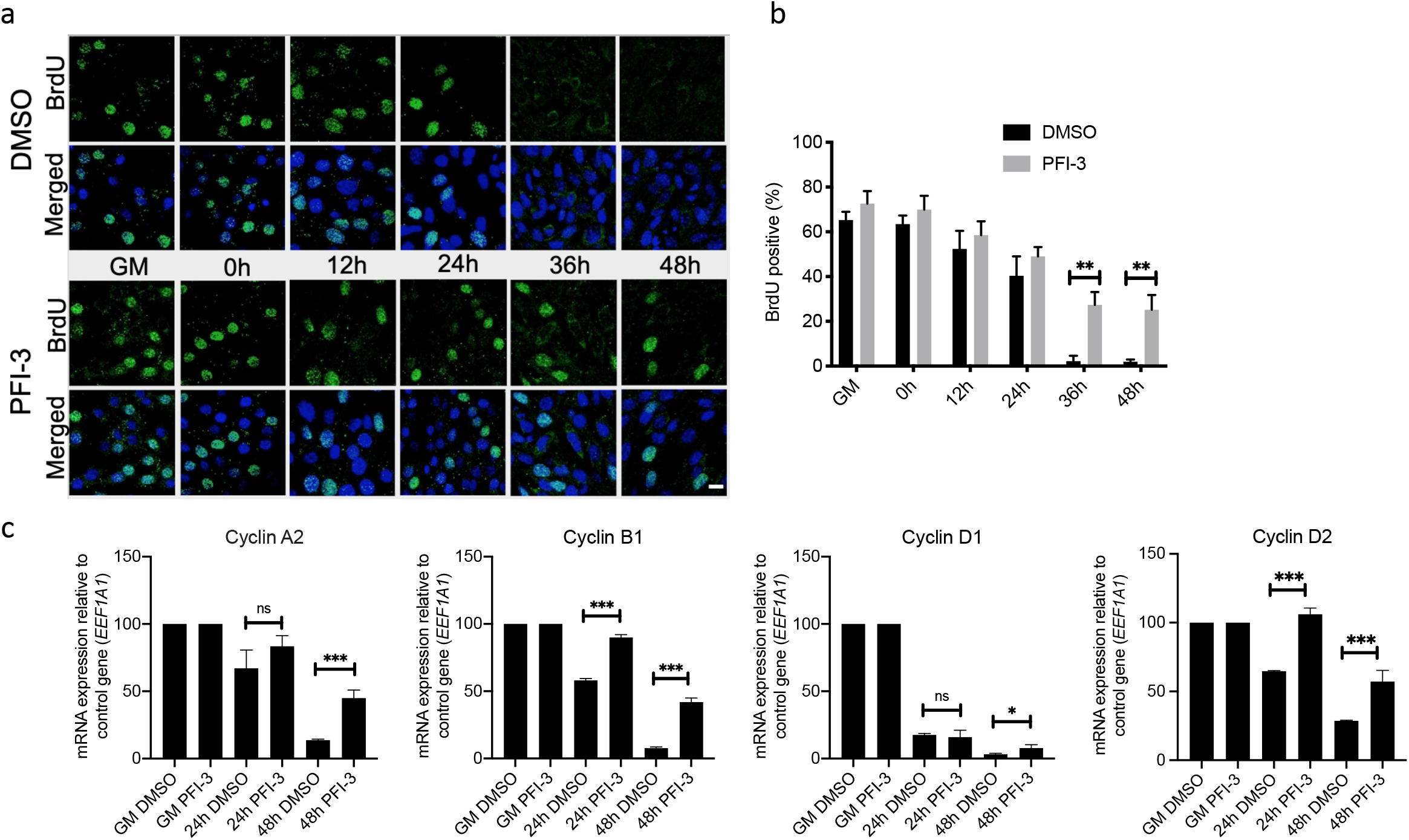
(a) C2C12 cells treated with PFI-3 show continued BrdU incorporation after 36h and 48 h post-induction of differentiation as compared to control cells, scale 5 μm. (b) Quantification of confocal images for BrdU incorporation assay in DMSO or PFI-3 treated C2C12 cells at the indicated timepoints. (c) mRNA expression levels of cyclin A2, cyclin B1, cyclin D1, and cyclin D2 in C2C12 myoblasts treated with DMSO or PFI-3 for indicated timepoints. Expression was normalized to a control gene (EEF1A1). 100% expression is defined as the timepoint at which maximal expression was observed. ns, not significant, *p<0.05, **p<0.01 and ***p<0.005

### BAF180 is dispensable for C2C12 myoblast differentiation

The composition of mSWI/SNF complexes is variable depending on function, cell-type and context. Every functional mSWI/SNF complex contains either the BRG1 or the BRM ATPase, while one major sub-class of mSWI/SNF complexes also contains BAF180 (10, 13). Thus, PFI-3 treatment affects all mSWI/SNF complexes. BRG1 and BRM have been shown to be required for skeletal muscle differentiation (25, 26, 28, 30, 56, 57), but the requirement for BAF180 in this process has not been evaluated.

We knocked down BAF180 using siRNA. C2C12 cells depleted for BAF180 were induced for differentiation alongside cells with scrambled siRNA treatment (Fig 5a, 5b). The cells showed no phenotypic defect and differentiated normally. This result suggests that BAF180 may be dispensable for myogenesis and implies that PFI-3 induced inhibition of myoblast differentiation is mediated through inactivation of BRG1 and/or BRM bromodomain function.

**Fig. 5.**
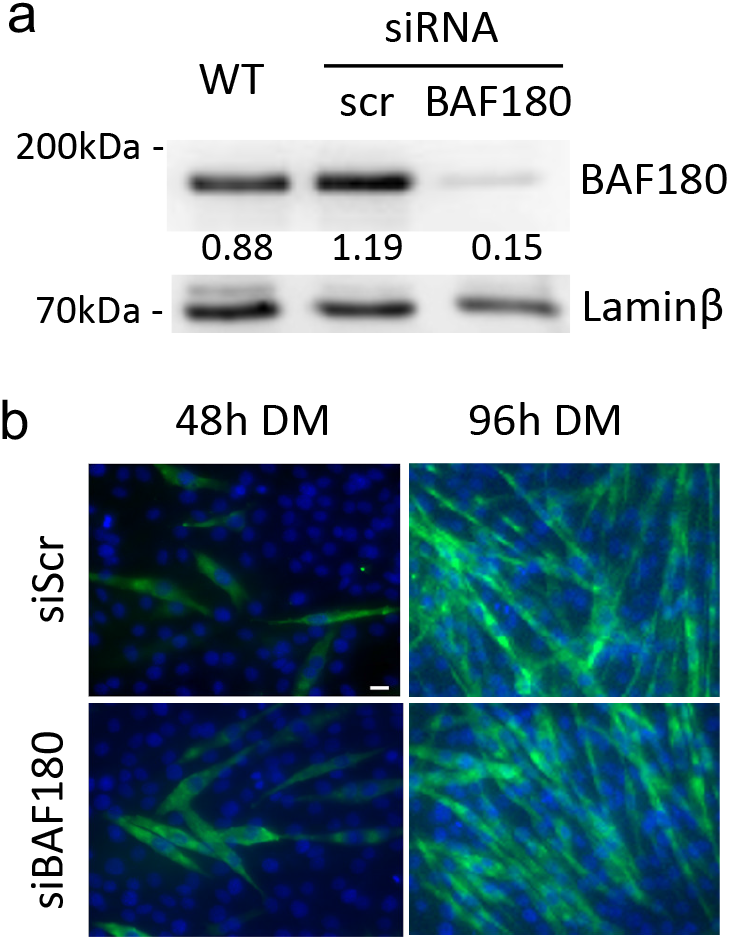
(a) Western blot analysis showed siRNA-mediated silencing of endogenous BAF180 in proliferating C2C12 cells. A scramble siRNA (siScr) was used as a control. The indicated numbers are the pixel counts normalized to Laminβ expression calculated using ImageJ. (b) Representative images of myosin heavy chain staining in 48h and 96h differentiated cells transfected with the control or BAF180-targeting siRNAs. The cells were fixed and analyzed by immunofluorescence using an anti-myosin heavy chain mAb MF20 (*green*). The nuclei were visualized by DAPI staining (*blue*). Scale bar, 20 μm.

### Gene targets of PFI-3 inhibition of mSWI/SNF bromodomains predominantly overlap with targets of BRG1 knockdown during myogenesis

The importance of BRG1 and BRM in skeletal muscle differentiation has been shown previously by multiple groups. These studies have looked at muscle-specific gene expression profiles and promoter binding of selected mSWI/SNF subunits on myogenic regulatory sequences. In a recently published study, the authors performed siRNA-mediated knockdown of BRG1 in C2C12 cells differentiated for 48h followed by RNA-sequencing analysis (58). We compared the siBRG1 dataset from this study with our RNA-sequencing dataset generated from PFI-3 treated C2C12 cells differentiated under similar conditions (Fig 6a). The rationale behind this comparison was to get an understanding of the relative importance of the BRG1 bromodomain. The analysis showed that 46% of the gene targets downregulated due to PFI-3 treatment overlapped with downregulated genes in the siBRG1 dataset (Fig 6a). Similarly, 46% of the upregulated genes due to PFI-3 inhibition were common with upregulated genes from siBRG1 dataset. The results show that a subset of BRG1-dependent gene expression in differentiating myoblasts requires bromodomain function. GO analysis of the common overlapping genes was conducted. Common downregulated genes belonged to muscle differentiation related processes while the common upregulated targets fell into cell-cycle related categories. (Fig 6a-b; Supp. Table 5). Promoters of the common upregulated and downregulated genes were also analyzed using HOMER to search for the presence of known consensus motifs within 1kb upstream of their TSS (Fig 6a-b; Supp. Table 5). Promoters of upregulated genes contained motifs known to be bound by E2F, KLF, NFY and Sp1 TF families while those of downregulated genes were enriched for motifs corresponding to MEF and MRF muscle-specific transcription factor families (Fig 6a-b; Supp. Table 5). Thus, the results from the GO and HOMER motif enrichment analyses of overlapping genes and their promoters are similar to those from PFI-3 treatment as shown in Fig 3c-d and therefore point towards a crucial role played by BRG1 bromodomain in skeletal muscle differentiation and cell cycle regulation.

**Fig. 6.**
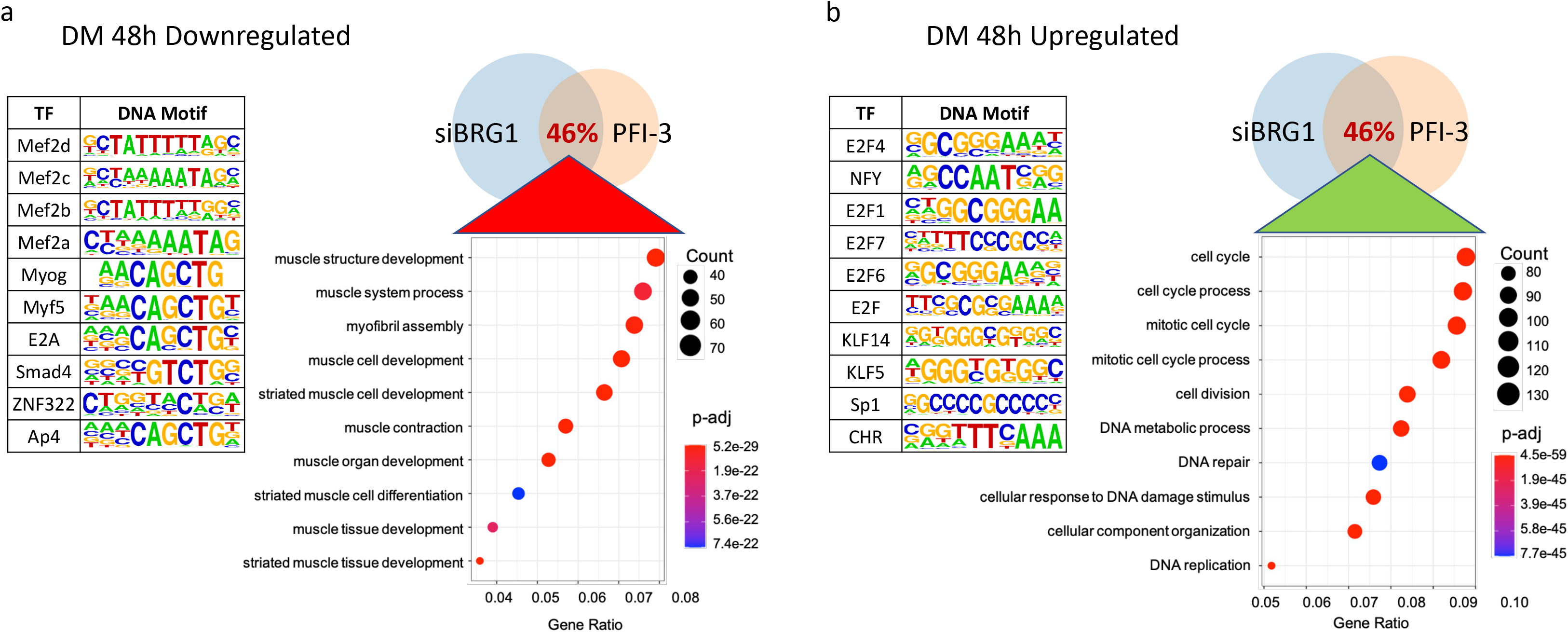
Comparative analysis of RNA-seq datasets from BRG1 knockdown performed by Zhu et al. (NAR, 2020) and from PFI-3 treatment. Venn diagrams represent DEGs in corresponding datasets. The overlap represents genes common to both datasets. GO and HOMER motif enrichment analyses show biological process categories and motifs identified within 1kb upstream of the TSS in the promoters of (a) common downregulated genes and (b) common upregulated genes.

We examined the genes that were dependent on BRG1 for expression but independent of PFI-3-mediated inhibiton of bromodomain function (Supp. Fig 3; Supp. Table 6). Genes that are up- and down-regulated predominantly represented targets involved in metabolic processes and do not include genes that control skeletal muscle differentiation or control of cell cycle. This suggests that bromodomain-dependent regulation of gene expression is critical for myogenesis. This result also is consistent with prior studies showing that ATPase domain function is required for BRG1-mediated regulation of metabolism (59, 60) and that PFI-3 treatment did not affect cancer cell proliferation dependent on BRG1 and/or BRM (40, 60).

Microarray analysis of gene expression in C2C12 cells upon siRNA-mediated knockdown of BRG1 or BRM has also been done in a prior study by Albini et. al (30). Despite the difference in methodologies, we overlapped the DEGs from that study with our RNA-seq data from PFI-3 treated C2C12 cells at comparable timepoints (48h post-differentiation) to evaluate if the outcome from this comparison is in consonance with the previous results. The analysis showed that more than one-third of the genes identified by Albini et al. as unique BRG1 targets overlapped with differentially expressed genes from our PFI-3 RNA-seq dataset (Supp Fig 4a). Upon looking individually at upregulated and downregulated targets uniquely regulated by BRG1, there was a 39% and 34% overlap, respectively (Supp Fig 4b). These common overlapping target genes were then characterized using GO analysis (Supp Fig 4a-b). Upregulated genes fell into cell-cycle related categories, and downregulated genes belonged to muscle differentiation related categories (complete analysis in Supp. Table 7). This is in agreement with the overall results from PFI-3 treatment indicating the importance of active bromodomain for BRG1 function. Of the genes identified by Albini et al. as unique BRM targets, only about 20% were common with PFI-3 DEGs (Supp Fig 4b). These numbers show that more genes may be regulated by the BRG1 bromodomain as compared to the BRM counterpart. Albini et al. also identified a set of genes that were coregulated by both BRG1 and BRM. About 32% of these upregulated targets and 16% of the downregulated targets were found in the corresponding list of differentially expressed genes due to PFI-3 induced bromodomain inhibition (Supp Fig 4b; Supp. Table 7). This comparative analysis further confirms that both BRG1 and BRM bromodomains play a role in skeletal myogenesis.

### PFI-3 inhibition of mSWI/SNF bromodomains decreased binding of BRG1 and BRM to target gene promoters

BRG1-containing mSWI/SNF complexes are recruited to regulatory regions of myogenic genes upon induction of muscle differentiation (25, 26, 29, 30, 61, 62). This step is required to induce remodeling of chromatin at myogenic gene loci, thus allowing activation of muscle-specific gene expression.

We performed ChIP experiments to determine whether the occupancy of BRG1 was affected at the myogenic gene regulatory regions in response to bromodomain inhibition by PFI-3. As expected, BRG1 occupancy at myogenic regulatory regions increased as a function of differentiation in DMSO-treated samples (Fig 7a). However, BRG1 occupancy at the tested regulatory sequences was partly inhibited in PFI-3 treated cells. This loss of binding correlates with the decreased expression of these genes as seen in previous results (Fig. 2). We also looked at binding of BRG1 and BRM on cyclin D1 and cyclin D2 promoters. It is known that in C2C12 myoblasts differentiated for 48h, BRM binding to the cyclin D1 gene promoter is crucial for its repression and cell-cycle exit (30). In that study, the authors showed that the expression of cyclin D1 was co-regulated by both BRG1 and BRM during the later stages of differentiation (30). In MCF-7 cells, BRG1 has been shown to bind to cyclin D1 promoter and regulate its expression (63). In PFI-3 treated C2C12 cells, chromatin IPs at the cyclin D1 promoter showed a decrease in occupancy of BRM and BRG1 (Fig 7b). Similarly, we saw a significant decrease in binding of BRG1 and BRM at the cyclin D2 promoter due to PFI-3 treatment. These results show that the bromodomain function of BRG1 and BRM contributes to their binding at target gene promoters.

**Fig. 7.**
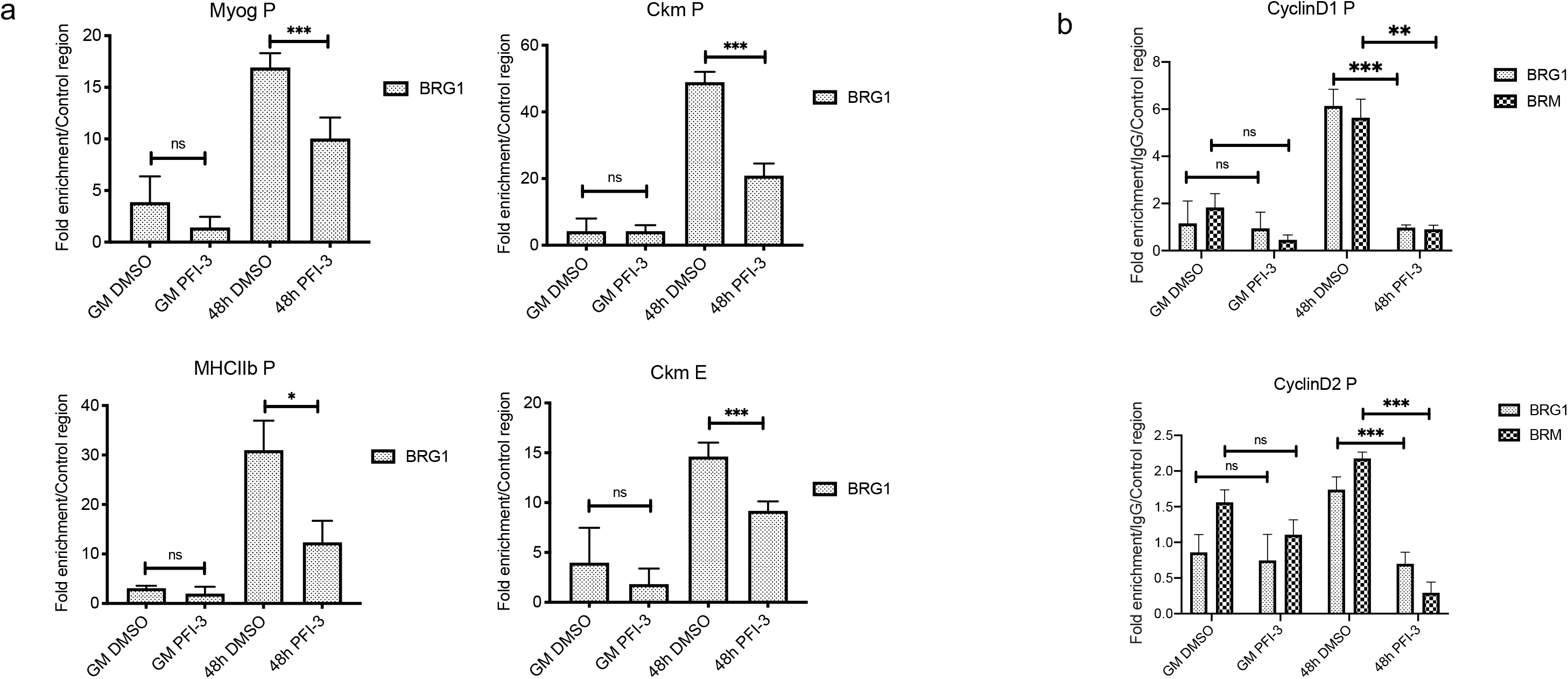
ChIP assays show decreased occupancy of BRG1 and BRM on regulatory regions of target genes upon PFI-3 inhibition. (a) Bar plots for BRG1 occupancy on the myogenin promoter (Myog P), myosin heavy chain promoter (MHCIIb P), creatine kinase promoter (Ckm P) and creatine kinase enhancer (Ckm E) are shown. (b) Bar plots for BRG1 and BRM occupancies on the cyclin D1 and cyclin D2 promoters. The values have been normalized to an IgG experimental control. These values are also normalized for binding at a non-specific region. ns, not significant, *p<0.05, **p<0.01 and ***p<0.005

**Fig. 8.**
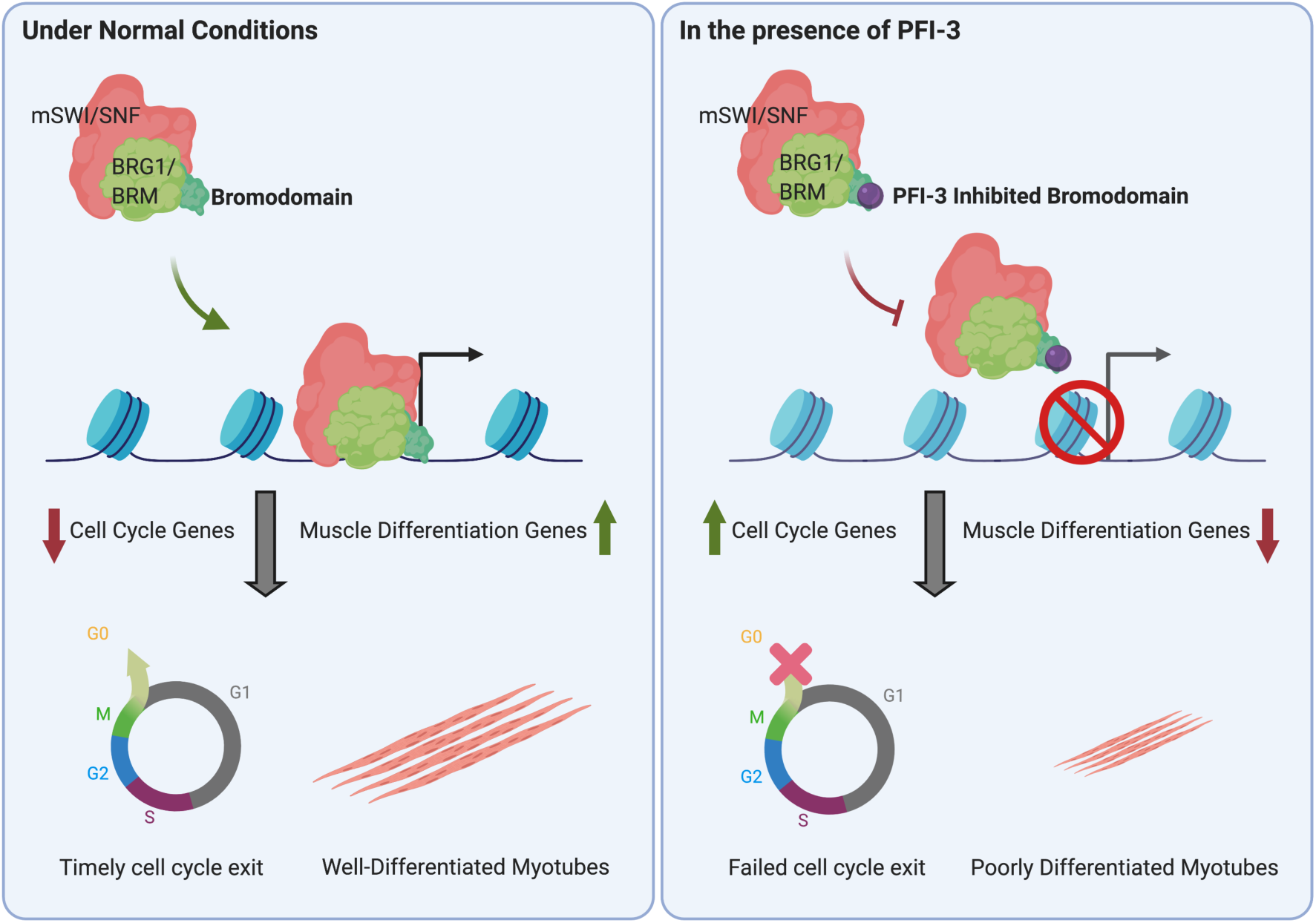
Graphical summary showing the effects of PFI-3 induced bromodomain inhibition on skeletal myogenesis. In normal conditions, BRG1/BRM with active bromodomains can bind to promoters of target genes when muscle differentiation is induced. This in turn affects two important aspects of skeletal myogenesis: cell cycle exit and the formation of differentiated multinucleated myotubes. In the presence of PFI-3, BRG1 and BRM show reduced binding to target gene promoters leading to continued cell-cycle and incomplete differentiation resulting in shorter myotubes.

## DISCUSSION

### Bromodomains in mSWI/SNF proteins

Bromodomains are a conserved structural motif found in only 46 human proteins, and they are classified into eight families (38). Bromodomains bind to acetylated lysines, which facilitates protein-protein interactions (37). The ability of bromodomains to target proteins to acetylated nucleosomes containing acetylated histones has been predicted to be a mechanism by which chromatin epigenetic modifications are read, thereby enabling translation of the histone mark via the bromodomain protein or via proteins associated with the bromodomain-containing protein (37, 64, 65). Family VIII bromodomains include the 6 bromodomains found in the N-terminal portion of the BAF180 protein, the bromodomains present in the BRG1 and BRM ATPases, and the bromodomain found in the histone lysine N-methyltransferase ASH1L (38).

Work to date on mSWI/SNF bromodomains has consisted of structural analyses (66–70), in vitro studies of bromodomain binding to histones, DNA and nucleosomes (67, 70–74) and a limited number of functional analyses in higher eukaryotes (75, 76). Of particular note, deletion of the bromodomain in *Drosophila* BRM, which is the only SWI/SNF ATPase, had no effect on developing or adult flies (77). In human cells, reconstitution of BRG1-deficient tumor cells with wildtype or mutant versions of BRG1 determined that sequences C-terminal to the ATPase domain, which includes the bromodomain, were not required for BRG1-mediated co-activation of transcription by the glucocorticoid receptor (78). Similarly, BRG1 containing a bromodomain mutation was capable of co-activating myocardin to promote smooth muscle-specific gene expression (79). However, the BRG1 bromodomain directly contributes to the ability of Repressor Element 1-silencing Transcription Factor (REST) to bind chromatin and repress target genes (80). Mutation of each of the six BAF180 bromodomains revealed that four of the six promoted tumor suppressor function, gene regulation, and chromatin affinity in clear cell renal cell carcinoma cells (81). Other work showed that the 4^th^ bromodomain of BAF180 mediated interaction with acetylated p53, which promotes p53 binding to and transcriptional activation of its target promoters (82). Thus, the requirement for functional mSWI/SNF protein bromodomains is variable. It may be cell-type dependent as well. Here we demonstrate that the BRG1 and/or BRM bromodomains, but not the BAF180 bromodomains, contribute to myogenic differentiation. Both BRG1 and BRM are required for myogenic differentiation; this work is the first to identify a contribution to myogenesis by any domain other than the ATPase domain. The work further supports the idea of context-dependent requirements for mSWI/SNF bromodomain functions.

### Use of PFI-3 to probe biological function of mSWI/SNF subunits containing bromodomains

Efforts to identify pharmacological inhibitors of bromodomains identified salicylic acid as a specific interactor of BRG1, BRM, and BAF180 bromodomains (41, 83). This led to a series of structure-guided design steps that resulted in the PFI-3 inhibitor that is specific for the second and fifth bromodomain of BAF180 and the bromodomains of BRG1 and BRM (41, 42).

To date PFI-3 has been used to probe biological function in a number of contexts. Because BRG1 and other subunits of the mSWI/SNF enzyme have been shown to be required for proliferation of some cancer cells (84–86), PFI-3 was tested for inhibitory effects on cancer cell proliferation, with no effect observed on many different cancer cell types, including the NCI-60 tumor cell panel (40, 41, 87). These results demonstrate that mSWI/SNF bromodomains are not required for cancer cell proliferation. An inhibitory effect of PFI-3 was observed in PTEN-depleted prostate cancer cells in culture, in xenografts and in PTEN deficient mouse model susceptible to prostate cancer (88). PTEN is a tumor suppressor that normally regulates the AKT/PKB signaling pathway (89). Thus, the importance of mSWI/SNF bromodomains, and specifically, the BRG1 bromodomain, is enhanced in the absence of PTEN and inhibition of AKT/PKB signaling, suggesting a complex mode of regulation of mSWI/SNF protein bromodomain function. PFI-3 treatment increased the accessibility of an mSWI/SNF-repressed promoter and its gene expression (90). In these and other studies, PFI-3 altered both gene expression patterns dependent on mSWI/SNF bromodomain-containing proteins and the cellular and organismal phenotypes controlled by those genes (87, 88, 91–96). The data indicate that the pleiotropic effects of PFI-3 and inhibition of mSWI/SNF bromodomains links to the ability of the chromatin remodeling enzyme to modulate gene expression.

In the realm of tissue specification, PFI-3 treatment caused a loss of “stemness” and promoted differentiation of ESCs and trophoblast and neural stem cells in the absence of differentiation signaling (41, 97). In other contexts, PFI-3 inhibited differentiation, blocking the ability of myoblasts and pre-adipocytes to form myotubes and adipocytes, respectively, in the presence of differentiation signaling (42). In this report, we investigated the mechanisms responsible for the inhibitory effects of PFI-3 on myogenesis. PFI-3 treatment affected bromodomain function of BRG1 and BRM, but BAF180, as BAF180 was dispensable for differentiation. BRG1/BRM bromodomain function was required for appropriate regulation of cell cycle withdrawal and initiation of the myogenic gene expression program, with mis-regulation of a subset of the genes regulated by BRG1 and BRM. Deficient gene regulation was linked to the partial inhibition of the ability of BRG1 and BRM to bind to target gene regulatory sequences, reflecting a necessary contribution of these bromodomains to promote interaction of the mSWI/SNF enzymes with chromatin.

### BAF180 is dispensable for myogenesis

mSWI/SNF complexes are a family of enzyme complexes marked by diversity of subunit composition (6, 11, 98). Initial descriptions of mSWI/SNF complexes reported two separable biochemical fractions that showed ATP-dependent chromatin remodeling activity (8–10). These complexes have become known as BAF (BRG1/BRM-associated factors) and PBAF (Polybromo-associated BAF), the latter taking its name from the presence of the BAF180 protein that is specific to this complex. However, both BAF and PBAF complexes themselves are merely separable groups of complexes that contain both shared and unique subunits (99, 100). A third family of complexes, called ncBAF (noncanonical BAF) was identified more recently (13, 101).

Although there are many subtypes of functional mSWI/SNF complexes in the cell, all of them necessarily have at least one of the proteins from the family VIII of bromodomains (102). BAF and ncBAF complexes contain either BRG1 or BRM, while PBAF complexes contain BRG1 and BAF180 (13). BAF180 has been implicated in DNA damage repair (103, 104) and is also required for cardiac development (105, 106); knockout in mice caused severe hypoplastic ventricle development and trophoblast placental defects (105). However, adult mice with BAF180 depletion were phenotypically normal except for a hematopoietic stem cell defect observed in aged mice (107). Similarly, BAF180 shows tumor-suppressive properties in some but not all cancer cell lines (107–111). These findings are consistent with context-specific requirements for the BAF180 protein.

In our work, we used siRNA-mediated knockdown to show that BAF180 is dispensable for skeletal muscle differentiation. This suggests that the PBAF family of mSWI/SNF complexes are also dispensable in this differentiation program. Although there are many reports characterizing the requirement for mSWI/SNF complexes in myogenesis, focus has been limited to the two ATPase subunits, to BAF47/INI1, which is shared by BAF and PBAF complexes, and to the BAF60 subunit that is shared by all subfamilies of mSWI/SNF complexes (25–27, 29–31, 56, 57, 112–125). A prior report documented the binding of BAF250A to myogenic promoters (115), perhaps implicating BAF complexes as the relevant mSWI/SNF enzyme subfamily for myogenic differentiation, but the requirement for BAF250A was not evaluated. Nevertheless, a requirement for specialized complexes for specific gene regulation events is one of the main hypotheses for existence of diverse families of mSWI/SNF complexes.

### BRG1 and BRM bromodomain function contribute to the regulation of myogenic differentiation

BRG1 and BRM contribute to the activation of the myogenic gene expression program and BRM contributes to the cell cycle arrest of myoblasts that precedes differentiation (25–30, 56). Inhibition of the mSWI/SNF bromodomains by PFI-3 recapitulated these findings, indicating that the bromodomains of BRG1 and BRM are needed to both regulate cell cycle exit and for the initiation of tissue-specific gene expression. RNA-seq analysis of PFI-3 treated cells provided evidence of global disruption of the regulation of cell cycle exit and the initiation of myogenic gene expression. A recently published RNA-seq study investigating the role of chromatin remodeling in skeletal myogenesis performed knockdown of BRG1 and evaluated gene expression at timepoints comparable with our study (58). Comparison of this dataset with ours identified a large overlapping subset of gene targets involved in cell cycle exit and myogenesis indicating that BRG1 bromodomain plays a crucial role in regulation of BRG1-dependent events in skeletal muscle differentiation.

Additionally, prior microarray-based studies of myogenic gene expression upon BRG1 and BRM knockdown also identified these processes as being BRG1- and BRM-dependent (30). Despite the differences in platforms, we integrated the two datasets. There was a considerably greater overlap between PFI-3 affected genes and genes mis-regulated by BRG1 knockdown than there was between PFI-3 affected genes and genes mis-regulated by BRM knockdown. This may suggest that a greater percentage of genes that require BRG1 are also dependent on the BRG1 bromodomain than is true for the set of genes that require BRM.

Regardless, our ChIP experiments demonstrated that bromodomain inhibition resulted in a decreased ability of BRG1 to bind to genes activated during the myogenic differentiation protocol and a decreased ability of BRG1 and BRM to bind to genes controlling cell cycle. The chromatin interacting properties of the BRG1 and BRM bromodomains therefore likely contribute to gene expression. The principles determining the variable requirements for BRG1, BRM and BAF180 bromodomains in different cellular contexts remain to be investigated. However, many of the mSWI/SNF subunits have additional domains that could promote chromatin interaction. For instance, BRG1 and BRM proteins have AT hooks, BAF180 and the BAF57 protein have an HMG box-like domain, the ARID1A/1B and ARID2 proteins have ARID domains that may mediate nucleic acid interactions, and BRD7 and BRD9 also have bromodomains. The requirement for any of these domains may be dependent on the protein makeup of the particular mSWI/SNF complex and the presence or absence of these other domains. Such a scenario suggests that these largely non-sequence specific DNA-binding domains function in an additive or cooperative manner to help facilitate chromatin interactions and remodeling events. Additional characterization of the putative chromatin interactions domains in the context of differentiation will be necessary to determine whether they are required and act in concert to promote mSWI/SNF interaction with chromatin and function.

## MATERIALS AND METHODS

### Antibodies and Chemicals

Antibodies were purchased from Santa Cruz Biotech, USA (anti-BRG1, sc-17796; anti-BrdU, sc-32323). Myosin Heavy chain antibody (#MF20) was purchased from the Developmental Studies Hybridoma Bank, University of Iowa, USA. BRM antisera was described previously (126). Lysis buffers for ChIP assays were purchased from Cell Signaling Technology, USA (SimpleChIP^®^ Enzymatic Cell Lysis Buffers A & B, 14282; SimpleChIP^®^ Chromatin IP Buffers, 14231). Dulbecco’s modified Eagle's medium (DMEM) was purchased from ThermoFisher Scientific (#11965118).

### Cell culture

C2C12 cells were purchased from ATCC (Manassas, VA) and maintained at sub-confluent densities in DMEM supplemented with 10% FBS and 1% penicillin/streptomycin in a humidified incubator at 37°C in 5% CO_2_.

Mouse satellite cells were isolated from leg muscles of 2 week old C57BL6/J mice using Percoll sedimentation followed by differential plating as described previously (114). Mice were housed in the animal care facility at the University of Massachusetts Medical School and used in accordance with a protocol approved by the Institutional Animal Care and Use Committee (IACUC).

For differentiation, cells at > 70% confluency were switched to DMEM medium supplemented with 2% horse serum and 2 μg/ml of bovine insulin (Sigma-Aldrich, St. Louis, MO). Where indicated, cells were pre-treated with DMSO or PFI-3 (Cayman Chemical, Ann Arbor, MI) for 24h before inducing differentiation. PFI-3 was maintained during the course of the experiment at 50μM and the medium was replaced every 24 hours.

### siRNA transfection

C2C12 cells were plated on 24-well plates in DMEM medium 24h before transfection. Cells were transfected at 30-40% confluence using the Lipofectamine 2000 (Invitrogen) reagent with 50 nM siRNA SMARTpool ON-TARGETplus (Dharmacon, Scrambled non-targeting Pool # D-001810-10-20 and Pbrm1 #L-044781-00-0005). 48h post-transfection, the cells were induced for differentiation and samples were harvested at indicated times for further analysis.

### Immunocytochemistry

Cells were seeded on 22mm x 22mm size coverslips in 35mm dishes and were harvested after the indicated treatments at the specified timepoints. The samples were washed with PBS twice and then fixed with ice-cold fixative (2% formaldehyde and 0.2% glutaraldehyde in PBS) for 10 minutes on ice. The cells were washed with PBS twice and permeabilized with ice-cold permeabilization buffer (0.2% Triton-X 100 in PBS) for 5 minutes on ice. Samples were then washed once with PBS and blocked using freshly prepared blocking reagent (5% BSA in PBS) for 30 minutes at room temperature (RT). The cells were washed twice with PBS and incubated with primary antibody cocktail diluted to the desired concentration in blocking reagent (2% BSA in PBS) for 2 hours at room temperature. Post-incubation, samples were washed thrice with PBS 10 minutes each to remove non-specific binding. Cells were then incubated with fluorophore-conjugated secondary antibody (1:100) and DAPI (2μg/ml), both diluted in blocking reagent (2% BSA in PBS) for 45 minutes at RT followed by 3 washes with PBS to remove non-specific staining. The stained cells on coverslips were then inverted-mounted on glass slides in 70% glycerol and the sides were sealed with nail paint. Confocal imaging analysis was performed using Leica TCS SP5 II laser scanning confocal microscope and analyzed with Leica Lite software.

### Fusion Index

For calculation of fusion index, cells were harvested at the specified timepoints after the indicated treatments and were immunostained with myosin heavy chain (MF20, DSHB) and DAPI (nuclear staining) as described above. The images were captured at 40X magnification using a Leica TCS SP5 II laser scanning confocal microscope. Analysis was performed by scoring cells for number of nuclei and MHC staining. Fusion index was calculated as the percentage of nuclei/cells stained with myosin heavy chain as compared to total number of nuclei/cells (43).

### RIPA buffer Lysis

Cells were harvested after the indicated treatments at specific timepoints and were washed twice with ice-cold PBS. After draining all residual PBS, cells were scraped into 1ml ice-cold PBS with 1X protease inhibitor cocktail (Sigma Aldrich, P8340) and pelleted at 1500 X *g* for 5 minutes at 4°C. The pellets were lysed in 500ul RIPA buffer (50 mM Tris-HCl, pH7.4, 150 mM NaCl, 1mM EDTA, 1% NP-40 and 0.25% sodium deoxycholate) supplemented with 1X protease inhibitor cocktail (Sigma Aldrich, P8340). Samples were incubated on ice for 30 minutes and whole cell extracts were prepared by passing the lysed pellets through a 27-gauge needle at least 4-5 times. Samples were centrifuged at 14000 X *g* for 10 minutes at 4°C and supernatants were collected.

### Western Blot Analysis

Protein concentrations were determined using a Pierce™BCA protein assay kit (ThermoFisher Scientific, USA) according to the manufacturer’s protocol. Equal amounts of protein from each sample were aliquoted and mixed with 4X Laemmli Sample Buffer (BioRad) and boiled at 95°C for 10 minutes. The samples were electrophoresed on denaturing SDS-polyacrylamide gels and transferred onto Immobilon-P PVDF membranes (Merck Millipore, USA). The membranes were then blocked using 5% non-fat milk in PBS for 30 minutes followed by overnight incubation at 4°C with primary antibody against protein of interest at the desired dilution in 2% non-fat milk prepared in PBS. This was followed by 3 washes with TBS containing 0.1% Tween-20 for 5 minutes each at room temperature. The membranes were then incubated with HRP-conjugated anti-mouse or anti-rabbit secondary antibodies (1:2500, GE Healthcare Life Sciences) diluted in 2% non-fat milk prepared in PBS for 1 hour at RT followed by 3 washes with TBS containing 0.1% Tween-20 for 5 minutes each at room temperature. Chemiluminescent detection was performed with ECL Plus (GE Healthcare Life Sciences) using an Amersham Imager 600 (GE Healthcare Life Sciences). Representative blots from 3 independent experiments are shown. Band signal intensities were quantified using ImageJ software (NIH) (127).

### RNA isolation and quantitative RT-PCR

For RNA isolation, cells were grown in 35mm dishes and harvested after the indicated treatments at specified timepoints. The media were removed, and cells were washed twice with PBS before adding 1ml of Trizol (ThermoFisher) to each sample. RNA extraction was performed as per the manufacturer’s protocol. The final RNA pellet was resuspended in 50μl nuclease-free water. RNA concentrations were quantified using a Nanodrop1000 spectrophotometer (ThermoFisher Scientific). cDNA was prepared using 2μg of total RNA using Superscript III First Strand Synthesis Kit (Invitrogen) according to manufacturer’s protocol.

For qRT-PCR, 15μl reactions were prepared in duplicate for all desired samples using 1μl each of forward and reverse primers (10μM stocks) and 2μl cDNA template, and the volume was brought to 7.5μl using UltraPure distilled water. 7.5μl of Fast SYBR Green 2X Master Mix (Applied Biosystems) was added to the reaction. The samples were run using the default protocol in QuantStudio 3 RT-PCR machine (Applied Biosystems). Fold-changes were calculated using the 2^−ΔΔCt^ method (128). Primer sequences are listed in Supp. Table 8.

### RNA-sequencing analysis

For RNA sequencing, RNA samples were prepared as described above. Duplicate samples for each timepoint were evaluated for quality and concentration at the UMass Medical School MBCL Fragment Analyzer services. QC-approved samples were sent to BGI Americas Corporation for library preparation and RNA sequencing (129). Libraries were sequenced using the BGISEQ-500 platform and reads were filtered to remove adaptor-polluted, low quality and high content of unknown base reads. About 99% of the raw reads were identified as clean reads (~65M). Of these, about 70% of the reads were uniquely mapped to mouse reference genome mm10 using HISAT (130). Transcripts were reconstructed using StringTie (131), and novel transcripts were identified using Cufflinks (132). All transcripts were then combined and mapped to the mm10 reference transcriptome using Bowtie2 (133). Gene expression levels were calculated using RSEM (134). DEseq2 (135) and PoissonDis (136) algorithms were used to detect differentially expressed genes (DEG). GO analysis was performed on DEGs to cluster genes into function-based and pathway-based categories (48, 49). Motif analysis was performed using HOMER motif discovery software as described previously (49). For each differentially expressed gene, motif enrichment analysis was performed at promoters for locations up to 1 kb upstream of the TSS.

### BrdU incorporation and immunostaining

Cell were grown on coverslips in 35mm dishes with the indicated treatments. The culture media was replaced with fresh media containing 10mM BrdU 30 minutes prior to harvesting to allow the cells in S-phase to incorporate the nucleoside analog. Harvested samples were fixed and permeabilized as described above for immunocytochemistry. For DNA hydrolysis, samples were then incubated with 1.2N HCl diluted in PBS for 1 hour at 37°C, followed by neutralization in 0.1M sodium borate for 5 minutes at RT. The samples were washed with PBS thrice and immunostained as mentioned above.

### Chromatin Immunoprecipitation assay

Chromatin immunoprecipitation assays were performed as described previously (115). Quantification was performed using the fold enrichment method (2^-(Ct sample – Ct IgG)^) and shown as relative to a control region. Sequences of primers used for ChIP assays are listed in Supp. Table 8.

### Statistical analysis

All quantitative data for gene expression and chromatin immunoprecipitation are shown as mean +/− the standard deviation of at least three independent biological replicates. In the fusion index and BrdU incorporation assays, two independent experiments were performed in duplicate. A minimum of 200 nuclei per sample were counted and the results were expressed as the mean of calculated fusion index +/− standard deviation. Statistical analyses were performed using Graphpad Prism8 Student’s t-test with two-tailed distribution and equal variance (Graphpad Prism Software Inc.). Significance is displayed with *p<0.05, **p<0.01 and ***p<0.005.

## Data Availability

The data from this RNA-seq study has been deposited in NCBI’s Gene Expression Omnibus and are accessible through GEO accession number GSE151218.

## ACKNOWLEDGEMENTS

We thank T Padilla-Benavides, S Syed, and J Dilworth for comments and suggestions. This work was funded by NIH grants GM56244 and GM136393 to ANI.

## REFERENCES

1. Allshire RC, Madhani HD. 2018. Ten principles of heterochromatin formation and function. Nat Rev Mol Cell Biol 19:229–244.

2. Janssen A, Colmenares SU, Karpen GH. 2018. Heterochromatin: Guardian of the Genome. Annu Rev Cell Dev Biol 34:265–288.

3. Hota SK, Bruneau BG. 2016. ATP-dependent chromatin remodeling during mammalian development. Development 143:2882–2897.

4. De La Serna IL, Ohkawa Y, Imbalzano AN. 2006. Chromatin remodelling in mammalian differentiation: Lessons from ATP-dependent remodellers. Nat Rev Genet 7:461–473.

5. Clapier CR, Cairns BR. 2009. The Biology of Chromatin Remodeling Complexes. Annu Rev Biochem 78:273–304.

6. Hargreaves DC, Crabtree GR. 2011. ATP-dependent chromatin remodeling: genetics, genomics and mechanisms. Cell Res 21:396–420.

7. Wu JI. 2012. Diverse functions of ATP-dependent chromatin remodeling complexes in development and cancer. Acta Biochim Biophys Sin 44:54–69.

8. Kwon H, Imbalzano AN, Khavari PA, Kingston RE, Green MR. 1994. Nucleosome disruption and enhancement of activator binding by a human SW1/SNF complex. Nature 370:477–481.

9. Imbalzano AN, Kwon H, Green MR, Kingston RE. 1994. Facilitated binding of TATA-binding protein to nucleosomal DNA. Nature 370:481–485.

10. Wang W, Côté J, Xue Y, Zhou S, Khavari PA, Biggar SR, Muchardt C, Kalpana G V., Goff SP, Yaniv M, Workman JL, Crabtree GR. 1996. Purification and biochemical heterogeneity of the mammalian SWI-SNF complex. EMBO J 15:5370–5382.

11. Wang W, Xue Y, Zhou S, Kuo A, Cairns BR, Crabtree GR. 1996. Diversity and specialization of mammalian SWI/SNF complexes. Genes Dev 10:2117–2130.

12. Wu JI, Lessard J, Olave IA, Qiu Z, Ghosh A, Graef IA, Crabtree GR. 2007. Regulation of Dendritic Development by Neuron-Specific Chromatin Remodeling Complexes. Neuron 56:94–108.

13. Mashtalir N, D’avino AR, Michel BC, Cassel SH, Ranish JA, Kadoch Correspondence C. 2018. Modular Organization and Assembly of SWI/SNF Family Chromatin Remodeling Complexes. Cell 175:1272–1288.e20.

14. Chal J, Pourquié O. 2017. Making muscle: skeletal myogenesis in vivo and in vitro. Development 144:2104–2122.

15. Comai G, Tajbakhsh S. 2014. Molecular and cellular regulation of skeletal myogenesis. Curr Top Dev Biol, 1st ed. 110:1–73.

16. Motohashi N, Asakura A. 2014. Muscle satellite cell heterogeneity and self-renewal. Front Cell Dev Biol 2:1–14.

17. Chang NC, Rudnicki MA. 2014. Satellite Cells: The Architects of Skeletal Muscle. Curr Top Dev Biol 107:161–181.

18. Sambasivan R, Tajbakhsh S. 2015. Adult skeletal muscle stem cells. Results Probl Cell Differ 56:191–213.

19. Hernández-Hernández JM, García-González EG, Brun CE, Rudnicki MA. 2017. The myogenic regulatory factors, determinants of muscle development, cell identity and regeneration. Semin Cell Dev Biol 72:10–18.

20. Moncaut N, Rigby PWJ, Carvajal JJ. 2013. Dial M(RF) for myogenesis. FEBS J 280:3980–3990.

21. Asfour HA, Allouh MZ, Said RS. 2018. Myogenic regulatory factors: The orchestrators of myogenesis after 30 years of discovery. Exp Biol Med 243:118–128.

22. Blackwell T, Weintraub H. 1990. Differences and similarities in DNA-binding preferences of MyoD and E2A protein complexes revealed by binding site selection. Science 250:1104–1110.

23. Taylor M V., Hughes SM. 2017. Mef2 and the skeletal muscle differentiation program. Semin Cell Dev Biol 72:33–44.

24. Pon JR, Marra MA. 2016. MEF2 transcription factors: Developmental regulators and emerging cancer genes. Oncotarget 7:2297–2312.

25. de la Serna IL, Ohkawa Y, Berkes C a, Bergstrom D a, Dacwag CS, Tapscott SJ, Imbalzano AN. 2005. MyoD targets chromatin remodeling complexes to the myogenin locus prior to forming a stable DNA-bound complex. Mol Cell Biol 25:3997–4009.

26. Ohkawa Y, Tachibana T, Imbalzano AN, Yoshimura S, Higashi C, Marfella CGA, Dacwag CS, Tachibana T, Imbalzano AN. 2007. Myogenin and the SWI / SNF ATPase Brg1 Maintain Myogenic Gene Expression at Different Stages of Skeletal Myogenesis *. J Biol Chem 282:6564–6570.

27. Simone C, Forcales SV, Hill DA, Imbalzano AN, Latella L, Puri PL. 2004. p38 pathway targets SWI-SNF chromatin-remodeling complex to muscle-specific loci. Nat Genet 36:738–743.

28. Ohkawa Y, Marfella CGA, Imbalzano AN. 2006. Skeletal muscle specification by myogenin and Mef2D via the SWI/SNF ATPase Brg1. EMBO J 25:490–501.

29. Forcales S V., Albini S, Giordani L, Malecova B, Cignolo L, Chernov A, Coutinho P, Saccone V, Consalvi S, Williams R, Wang K, Wu Z, Baranovskaya S, Miller A, Dilworth FJ, Puri PL. 2012. Signal-dependent incorporation of MyoD-BAF60c into Brg1-based SWI/SNF chromatin-remodelling complex. EMBO J 31:301–316.

30. Albini S, Toto PC, Dall’agnese A, Malecova B, Cenciarelli C, Felsani A, Caruso M, Bultman SJ, Puri PL. 2015. Brahma is required for cell cycle arrest and late muscle gene expression during skeletal myogenesis. EMBO Rep 16:1037–1050.

31. de la Serna IL, Roy K, Carlson KA, Imbalzano AN. 2001. MyoD can induce cell cycle arrest but not muscle differentiation in the presence of dominant negative SWI/SNF chromatin remodeling enzymes. J Biol Chem 276:41486–91.

32. Khavari PA, Peterson CL, Tamkunt JW, Mendel DB, Crabtree GR. 1993. BRG1 contains a conserved domain of the SWI2 / SNF2 family necessary for normal mitotic growth and transcription. Lett to Nat 366:170–174.

33. Muchardt C, Yaniv M. 1993. A human homologue of Saccharomyces cerevisiae SNF2/SWI2 and Drosophila brm genes potentiates transcriptional activation by the glucocorticoid receptor. EMBO J 12:4279–4290.

34. Chiba H, Muramatsul M, Nomoto A, Kato H. 1994. Two human homologues of Saccharomyces cerevisiae SWI2ISNF2 and Drosophila brahma are transcriptional coactivators cooperating with the estrogen receptor and the retinoic acid receptor. Nucleic Acids Res 22:1815–1820.

35. Meslamani J, Smith SG, Sanchez R, Zhou M-M. 2016. Structural features and inhibitors of bromodomains. Drug Discov Today Technol 19:3–15.

36. Moustakim M, Clark PGK, Hay DA, Dixon DJ, Brennan PE. 2016. Chemical probes and inhibitors of bromodomains outside the BET family. Medchemcomm 7:2246–2264.

37. Fujisawa T, Filippakopoulos P. 2017. Functions of bromodomain-containing proteins and their roles in homeostasis and cancer. Nat Rev Mol Cell Biol 18:246–262.

38. Filippakopoulos P, Picaud S, Mangos M, Keates T, Lambert JP, Barsyte-Lovejoy D, Felletar I, Volkmer R, Müller S, Pawson T, Gingras AC, Arrowsmith CH, Knapp S. 2012. Histone recognition and large-scale structural analysis of the human bromodomain family. Cell 149:214–231.

39. Xue Y, Canman JC, Lee CS, Nie Z, Yang D, Moreno GT, Young MK, Salmon ED, Wang W. 2000. The human SWI/SNF-B chromatin-remodeling complex is related to yeast Rsc and localizes at kinetochores of mitotic chromosomes. Proc Natl Acad Sci U S A 97:13015–13020.

40. Vangamudi B, Paul TA, Shah PK, Kost-Alimova M, Nottebaum L, Shi X, Zhan Y, Leo E, Mahadeshwar HS, Protopopov A, Futreal A, Tieu TN, Peoples M, Heffernan TP, Marszalek JR, Toniatti C, Petrocchi A, Verhelle D, Owen DR, Draetta G, Jones P, Palmer WS, Sharma S, Andersen JN. 2015. The SMARCA2/4 ATPase Domain Surpasses the Bromodomain as a Drug Target in SWI/SNF-Mutant Cancers: Insights from cDNA Rescue and PFI-3 Inhibitor Studies. Cancer Res 75:3865–3878.

41. Fedorov O, Castex J, Tallant C, Owen DR, Martin S, Aldeghi M, Monteiro O, Filippakopoulos P, Picaud S, Trzupek JD, Gerstenberger BS, Bountra C, Willmann D, Wells C, Philpott M, Rogers C, Biggin PC, Brennan PE, Bunnage ME, Schüle R, Günther T, Knapp S, Müller S. 2015. Selective targeting of the BRG/PB1 bromodomains impairs embryonic and trophoblast stem cell maintenance. Sci Adv 1:1–11.

42. Gerstenberger BS, Trzupek JD, Tallant C, Fedorov O, Filippakopoulos P, Brennan PE, Fedele V, Martin S, Picaud S, Rogers C, Parikh M, Taylor A, Samas B, O’Mahony A, Berg E, Pallares G, Torrey AD, Treiber DK, Samardjiev IJ, Nasipak BT, Padilla-Benavides T, Wu Q, Imbalzano AN, Nickerson JA, Bunnage ME, Müller S, Knapp S, Owen DR, Mahony AO, Berg E, Pallares G, Torrey AD, Treiber DK, Samardjiev IJ, Nasipak BT, Padilla-Benavides T, Wu Q, Imbalzano AN, Nickerson A, Bunnage ME, Mu S, Knapp S, Owen DR. 2016. Identification of a Chemical Probe for Family VIII Bromodomains through Optimization of a Fragment Hit. J Med Chem 59:4800–4811.

43. Metzinger L, Passaquin AC, Warter JM, Poindron P. 1993. α-Methylprednisolone promotes skeletal myogenesis in dystrophin-deficient and control mouse cultures. Neurosci Lett 155:171–174.

44. Millay DP, O’Rourke JR, Sutherland LB, Bezprozvannaya S, Shelton JM, Bassel-Duby R, Olson EN. 2013. Myomaker is a membrane activator of myoblast fusion and muscle formation. Nature 499:301–5.

45. Leikina E, Gamage DG, Prasad V, Kozlov MM, Chernomordik L V, Millay Correspondence DP, Goykhberg J, Crowe M, Diao J, Kozlov MM, Chernomordik L V, Millay DP. 2018. Myomaker and Myomerger Work Independently to Control Distinct Steps of Membrane Remodeling during Myoblast Fusion. Dev Cell 46:767–780.

46. Bi P, Ramirez-Martinez A, Li H, Cannavino J, McAnally JR, Shelton JM, Sánchez-Ortiz E, Bassel-Duby R, Olson EN. 2017. Control of muscle formation by the fusogenic micropeptide myomixer. Science 356:323–327.

47. Zhang Q, Vashisht AA, O’Rourke J, Corbel SY, Moran R, Romero A, Miraglia L, Zhang J, Durrant E, Schmedt C, Sampath SC, Sampath SC. 2017. The microprotein Minion controls cell fusion and muscle formation. Nat Commun 8:1–15.

48. Wei Huang D, Sherman BT, Lempicki RA. 2008. Bioinformatics enrichment tools: paths toward the comprehensive functional analysis of large gene lists. Nucleic Acids Res 37:1–13.

49. Heinz S, Benner C, Spann N, Bertolino E, Lin YC, Laslo P, Cheng JX, Murre C, Singh H, Glass CK. 2010. Simple Combinations of Lineage-Determining Transcription Factors Prime cis-Regulatory Elements Required for Macrophage and B Cell Identities. Mol Cell 38:576–589.

50. Müller H, Helin K. 2000. The E2F transcription factors: Key regulators of cell proliferation. Biochim Biophys Acta - Rev Cancer 1470:1–12.

51. Helin K. 1998. Regulation of cell proliferation by the E2F transcription factors. Curr Opin Genet Dev 8:28–35.

52. Dong JT, Chen C. 2009. Essential role of KLF5 transcription factor in cell proliferation and differentiation and its implications for human diseases. Cell Mol Life Sci 66:2691–2706.

53. Soufi A, Dalton S. 2016. Cycling through developmental decisions: How cell cycle dynamics control pluripotency, differentiation and reprogramming. Dev 143:4301–4311.

54. Id YM, Mckay Id DJ, Buttitta Id L. 2019. Changes in chromatin accessibility ensure robust cell cycle exit in terminally differentiated cells. Plos Biol 17:1–29.

55. Politis PK, Thomaidou D, Matsas R. 2008. Coordination of cell cycle exit and differentiation of neuronal progenitors. Cell Cycle 7:691–697.

56. de la Serna IL, Carlson KA, Imbalzano AN. 2001. Mammalian SWI/SNF complexes promote MyoD-mediated muscle differentiation. Nat Genet 27:187–190.

57. Roy K, de la Serna IL, Imbalzano AN. 2002. The myogenic basic helix-loop-helix family of transcription factors shows similar requirements for SWI/SNF chromatin remodeling enzymes during muscle differentiation in culture. J Biol Chem 277:33818–24.

58. Zhu X, Lan B, Yi X, He C, Dang L, Zhou X, Lu Y, Sun Y, Liu Z, Bai X, Zhang K, Li B, Li MJ, Chen Y, Zhang L. 2020. HRP2-DPF3a-BAF complex coordinates histone modification and chromatin remodeling to regulate myogenic gene transcription. Nucleic Acids Res 48:6563–6582.

59. Barutcu AR, Lajoie BR, Fritz AJ, McCord RP, Nickerson JA, Van Wijnen AJ, Lian JB, Stein JL, Dekker J, Stein GS, Imbalzano AN. 2016. SMARCA4 regulates gene expression and higherorder chromatin structure in proliferating mammary epithelial cells. Genome Res 26:1188–1201.

60. Wu Q, Madany P, Dobson JR, Schnabl JM, Sharma S, Smith TC, van Wijnen AJ, Stein JL, Lian JB, Stein GS, Muthuswami R, Imbalzano AN, Nickerson JA. 2016. The BRG1 chromatin remodeling enzyme links cancer cell metabolism and proliferation. Oncotarget 7:38270–38281.

61. Harada A, Okada S, Konno D, Odawara J, Yoshimi T, Yoshimura S, Kumamaru H, Saiwai H, Tsubota T, Kurumizaka H, Akashi K, Tachibana T, Imbalzano AN, Ohkawa Y. 2012. Chd2 interacts with H3.3 to determine myogenic cell fate. EMBO J 31:2994–3007.

62. Ohkawa Y, Mallappa C, Vallaster CSD, Imbalzano AN. 2012. Isolation of nuclei from skeletal muscle satellite cells and myofibers for use in chromatin immunoprecipitation assays. Methods Mol Biol 798:517–530.

63. Rao M, Casimiro MC, Lisanti MP, Amico MD’, Wang C, Shirley LA, Leader JE, Liu M, Stallcup M, Engel DA, Murphy DJ, Pestell RG. 2008. Inhibition of cyclin D1 gene transcription by Brg-1. Cell Cycle 7:647–655.

64. Haynes SR, Dollard C, Winston F, Beck S, Trowsdale J, Dawid IB. 1992. The bromodomain: a conserved sequence found in human, Drosophila and yeast proteins. Nucleic Acids Res 20:246–257.

65. Dhalluin C, Carlson JE, Zeng L, He C, Aggarwal AK, Zhou MM. 1999. Structure and ligand of a histone acetyltransferase bromodomain. Nature 399:491–496.

66. Singh M, Popowicz GM, Krajewski M, Holak TA. 2007. Structural ramification for acetyl-lysine recognition by the bromodomain of human BRG1 protein, a central ATPase of the SWI/SNF remodeling complex. ChemBioChem 8:1308–1316.

67. Singh M, D’Silva L, Holak TA. 2006. DNA-binding properties of the recombinant high-mobility-group-like AT-hook-containing region from human BRG1 protein. Biol Chem 387:1469–1478.

68. Chandrasekaran R, Thompson M. 2007. Polybromo-1-bromodomains bind histone H3 at specific acetyl-lysine positions. Biochem Biophys Res Commun 355:661–666.

69. Shen W, Xu C, Huang W, Zhang J, Carlson JE, Tu X, Wu J, Shi Y. 2007. Solution Structure of Human Brg1 Bromodomain and Its Specific Binding to Acetylated Histone Tails. Biochemistry 46:2100–2110.

70. Charlop-Powers Z, Zeng L, Zhang Q, Zhou MM. 2010. Structural insights into selective histone H3 recognition by the human Polybromo bromodomain 2. Cell Res 20:529–538.

71. Liao L, Alicea-Velázquez NL, Langbein L, Niu X, Cai W, Cho EA, Zhang M, Greer CB, Yan Q, Cosgrove MS, Yang H. 2019. High affinity binding of H3K14ac through collaboration of bromodomains 2, 4 and 5 is critical for the molecular and tumor suppressor functions of PBRM1. Mol Oncol 13:811–828.

72. Sanchez JC, Zhang L, Evoli S, Schnicker NJ, Nunez-Hernandez M, Yu L, Wereszczynski J, Pufall MA, Musselman CA. 2020. The molecular basis of selective DNA binding by the BRG1 AT-hook and bromodomain. Biochim Biophys Acta - Gene Regul Mech 1863:1–14.

73. Morrison EA, Sanchez JC, Ronan JL, Farrell DP, Varzavand K, Johnson JK, Gu BX, Crabtree GR, Musselman CA. 2017. DNA binding drives the association of BRG1/hBRM bromodomains with nucleosomes. Nat Commun 8:1–14.

74. Slaughter MJ, Shanle EK, McFadden AW, Hollis ES, Suttle LE, Strahl BD, Davis IJ. 2018. PBRM1 bromodomains variably influence nucleosome interactions and cellular function. J Biol Chem 293:13592–13603.

75. Wang X, Wang S, Troisi EC, Howard TP, Haswell JR, Wolf BK, Hawk WH, Ramos P, Oberlick EM, Tzvetkov EP, Vazquez F, Hahn WC, Park PJ, Roberts CWM. 2019. BRD9 defines a SWI/SNF sub-complex and constitutes a specific vulnerability in malignant rhabdoid tumors. Nat Commun 10:1–11.

76. Dutta A, Gogol M, Kim JH, Smolle M, Venkatesh S, Gilmore J, Florens L, Washburn MP, Workman JL. 2014. Swi/Snf dynamics on stress-responsive genes is governed by competitive bromodomain interactions. Genes Dev 28:2314–2330.

77. Elfring LK, Daniel C, Papoulas O, Deuring R, Sarte M, Moseley S, Beek SJ, Ross Waldrip W, Daubresse G, Depace A, Kennison JA, Tamkun JW. 1998. Genetic Analysis of brahma : The Drosophila Homolog of the Yeast Chromatin Remodeling Factor SWI2/SNF2. Genetics 148:251–265.

78. Trotter KW, Fan H-Y, Ivey ML, Kingston RE, Archer TK. 2008. The HSA Domain of BRG1 Mediates Critical Interactions Required for Glucocorticoid Receptor-Dependent Transcriptional Activation In Vivo. Mol Cell Biol 28:1413–1426.

79. Zhou J, Zhang M, Fang H, El-Mounayri O, Rodenberg JM, Imbalzano AN, Herring BP. 2009. The SWI/SNF chromatin remodeling complex regulates myocardin-induced smooth muscle-specific gene expression. Arterioscler Thromb Vasc Biol 29:921–8.

80. Ooi L, Belyaev ND, Miyake K, Wood IC, Buckley NJ. 2006. BRG1 chromatin remodeling activity is required for efficient chromatin binding by repressor element 1-silencing transcription factor (REST) and facilitates REST-mediated repression. J Biol Chem 281:38974–38980.

81. Porter EG, Dykhuizen EC. 2017. Individual bromodomains of Polybromo-1 contribute to chromatin association and tumor suppression in clear cell renal carcinoma. J Biol Chem 292:2601–2610.

82. Cai W, Su L, Liao L, Liu ZZ, Langbein L, Dulaimi E, Testa JR, Uzzo RG, Zhong Z, Jiang W, Yan Q, Zhang Q, Yang H. 2019. PBRM1 acts as a p53 lysine-acetylation reader to suppress renal tumor growth. Nat Commun 10:1–15.

83. Fedorov O, Niesen FH, Knapp S. 2012. Kinase Inhibitor Selectivity Profiling Using Differential Scanning Fluorimetry. Methods Mol Biol 795:109–118.

84. Baylin SB, Jones PA. 2016. Epigenetic Determinants of Cancer. Cold Spring Harb Perspect Biol 8:1–35.

85. Feinberg AP, Tycko B. 2004. The history of cancer epigenetics. Nat Rev Cancer 4:143–153.

86. Zaidi SK, Van Wijnen AJ, Lian JB, Stein JL, Stein GS. 2013. Targeting deregulated epigenetic control in cancer. J Cell Physiol 228:2103–2108.

87. Wu Q, Sharma S, Cui H, LeBlanc SE, Zhang H, Muthuswami R, Nickerson JA, Imbalzano AN. 2016. Targeting the chromatin remodeling enzyme BRG1 increases the efficacy of chemotherapy drugs in breast cancer cells. Oncotarget 7:27158–75.

88. Ding Y, Li N, Dong B, Guo W, Wei H, Chen Q, Yuan H, Han Y, Chang H, Kan S, Wang X, Pan Q, Wu P, Peng C, Qiu T, Li Q, Gao D, Xue W, Qin J. 2019. Chromatin remodeling ATPase BRG1 and PTEN are synthetic lethal in prostate cancer. J Clin Invest 129:759–773.

89. Georgescu MM. 2010. Pten tumor suppressor network in PI3K-Akt pathway control. Genes and Cancer 1:1170–1177.

90. Gillis NE, Taber TH, Bolf EL, Beaudet CM, Tomczak JA, White JH, Stein JL, Stein GS, Lian JB, Frietze S, Carr FE. 2018. Thyroid Hormone Receptor β Suppression of RUNX2 Is Mediated by Brahma-Related Gene 1–Dependent Chromatin Remodeling. Endocrinology 159:2484–2494.

91. Li Z, Xia J, Fang M, Xu Y. 2019. Epigenetic regulation of lung cancer cell proliferation and migration by the chromatin remodeling protein BRG1. Oncogenesis 8:1–14.

92. Minderjahn J, Schmidt A, Fuchs A, Schill R, Raithel J, Babina M, Schmidl C, Gebhard C, Schmidhofer S, Mendes K, Ratermann A, Glatz D, Nützel M, Edinger M, Hoffman P, Spang R, Längst G, Imhof A, Rehli M. 2020. Mechanisms governing the pioneering and redistribution capabilities of the non-classical pioneer PU.1. Nat Commun 11:1–16.

93. Li H, Lan J, Han C, Guo K, Wang G, Hu J, Gong J, Luo X, Cao Z. 2018. Brg1 promotes liver fibrosis via activation of hepatic stellate cells. Exp Cell Res 364:191–197.

94. Li Z, Lv F, Dai C, Wang Q, Jiang C, Fang M, Xu Y. 2019. Activation of Galectin-3 (LGALS3) Transcription by Injurious Stimuli in the Liver Is Commonly Mediated by BRG1. Front Cell Dev Biol 7:1–13.

95. Wang Z-J, Martin JA, Mueller LE, Caccamise A, Werner CT, Neve RL, Gancarz AM, Li J-X, Dietz DM. 2016. BRG1 in the Nucleus Accumbens Regulates Cocaine-Seeking Behavior-Supplementary info 80:652–660.

96. Ganguly D, Sims M, Cai C, Fan M, Pfeffer LM. 2018. Chromatin Remodeling Factor BRG1 Regulates Stemness and Chemosensitivity of Glioma Initiating Cells. Stem Cells 36:1804–1815.

97. Güneş C, Paszkowski-Rogacz M, Rahmig S, Khattak S, Camgöz A, Wermke M, Dahl A, Bornhäuser M, Waskow C, Buchholz F. 2019. Comparative RNAi Screens in Isogenic Human Stem Cells Reveal SMARCA4 as a Differential Regulator. Stem cell reports 12:1084–1098.

98. Kadoch C, Crabtree GR. 2015. Mammalian SWI/SNF chromatin remodeling complexes and cancer: Mechanistic insights gained from human genomics. Sci Adv 1:1–17.

99. Pulice JL, Kadoch C. 2016. Composition and function of mammalian SWI/SNF chromatin remodeling complexes in human disease. Cold Spring Harb Symp Quant Biol 81:53–60.

100. Hodges C, Kirkland JG, Crabtree GR. 2016. The many roles of BAF (mSWI/SNF) and PBAF complexes in cancer. Cold Spring Harb Perspect Med 6:1–25.

101. Michel BC, D’Avino AR, Cassel SH, Mashtalir N, McKenzie ZM, McBride MJ, Valencia AM, Zhou Q, Bocker M, Soares LMM, Pan J, Remillard DI, Lareau CA, Zullow HJ, Fortoul N, Gray NS, Bradner JE, Chan HM, Kadoch C. 2018. A non-canonical SWI/SNF complex is a synthetic lethal target in cancers driven by BAF complex perturbation. Nat Cell Biol 20:1410–1420.

102. Tang L, Nogales E, Ciferri C. 2010. Structure and function of SWI/SNF chromatin remodeling complexes and mechanistic implications for transcription. Prog Biophys Mol Biol 102:122–128.

103. Brownlee PM, Chambers AL, Cloney R, Bianchi A, Downs JA. 2014. BAF180 Promotes Cohesion and Prevents Genome Instability and Aneuploidy. Cell Rep 6:973–981.

104. Kakarougkas A, Ismail A, Chambers AL, Riballo E, Herbert AD, Künzel J, Löbrich M, Jeggo PA, Downs JA. 2014. Requirement for PBAF in Transcriptional Repression and Repair at DNA Breaks in Actively Transcribed Regions of Chromatin. Mol Cell 55:723–732.

105. Wang Z, Zhai W, Richardson JA, Olson EN, Meneses JJ, Firpo MT, Kang C, Skarnes WC, Tjian R. 2004. Polybromo protein BAF180 functions in mammalian cardiac chamber maturation. Genes Dev 18:3106–3116.

106. Huang X, Gao X, Diaz-Trelles R, Ruiz-Lozano P, Wang Z. 2008. Coronary development is regulated by ATP-dependent SWI/SNF chromatin remodeling component BAF180. Dev Biol 319:258–266.

107. Lee H, Dai F, Zhuang L, Xiao ZD, Kim J, Zhang Y, Ma L, You MJ, Wang Z, Gan B. 2016. BAF180 regulates cellular senescence and hematopoietic stem cell homeostasis through p21. Oncotarget 7:19134–19146.

108. Chowdhury B, Porter EG, Stewart JC, Ferreira CR, Schipma MJ, Dykhuizen EC. 2016. PBRM1 regulates the expression of genes involved in metabolism and cell adhesion in renal clear cell carcinoma. PLoS One 11:1–20.

109. Huang L, Peng Y, Zhong G, Xie W, Dong W, Wang B, Chen X, Gu P, He W, Wu S, Lin T, Huang J. 2015. PBRM1 suppresses bladder cancer by cyclin B1 induced cell cycle arrest. Oncotarget 6:16366–16378.

110. Xia W, Nagase S, Montia AG, Kalachikov SM, Keniry M, Su T, Memeo L, Hibshoosh H, Parsons R. 2008. BAF180 is a critical regulator of p21 induction and a tumor suppressor mutated in breast cancer. Cancer Res 68:1667–1674.

111. Gao W, Li W, Xiao T, Liu XS, Kaelin WG. 2017. Inactivation of the PBRM1 tumor suppressor gene amplifies the HIF-response in VHL−/− clear cell renal carcinoma. Proc Natl Acad Sci U S A 114:1027–1032.

112. Albini S, Coutinho P, Malecova B, Giordani L, Savchenko A, Forcales SV, Puri PL. 2013. Epigenetic Reprogramming of Human Embryonic Stem Cells into Skeletal Muscle Cells and Generation of Contractile Myospheres. Cell Rep 3:661–670.

113. Padilla-Benavides T, Nasipak BT, Imbalzano AN. 2015. Brg1 Controls the Expression of Pax7 to Promote Viability and Proliferation of Mouse Primary Myoblasts. J Cell Physiol 230:2990–2997.

114. Nasipak BT, Padilla-Benavides T, Green KM, Leszyk JD, Mao W, Konda S, Sif S, Shaffer S a., Ohkawa Y, Imbalzano AN. 2015. Opposing calcium-dependent signalling pathways control skeletal muscle differentiation by regulating a chromatin remodelling enzyme. Nat Commun 6:1–12.

115. Witwicka H, Nogami J, Syed SA, Maehara K, Padilla-Benavides T, Ohkawa Y, Imbalzano AN. 2019. Calcineurin Broadly Regulates the Initiation of Skeletal Muscle-Specific Gene Expression by Binding Target Promoters and Facilitating the Interaction of the SWI/SNF Chromatin Remodeling Enzyme. Mol Cell Biol 39:1–21.

116. Ochi H, Hans S, Westerfield M. 2008. Smarcd3 regulates the timing of zebrafish myogenesis onset. J Biol Chem 283:3529–3536.

117. Lickert H, Takeuchi JK, Von Both I, Walls JR, McAuliffe F, Adamson SL, Henkelman RM, Wrana JL, Rossant J, Bruneau BG. 2004. Baf60c is essential for function of BAF chromatin remodelling complexes in heart development. Nature 432:107–112.

118. Harada A, Mallappa C, Okada S, Butler JT, Baker SP, Lawrence JB, Ohkawa Y, Imbalzano AN. 2015. Spatial re-organization of myogenic regulatory sequences temporally controls gene expression. Nucleic Acids Res 43:2008–2021.

119. Padilla-Benavides T, Nasipak BT, Paskavitz AL, Haokip DT, Schnabl JM, Nickerson JA, Imbalzano AN. 2017. Casein kinase 2-mediated phosphorylation of Brahma-related gene 1 controls myoblast proliferation and contributes to SWI/SNF complex composition. J Biol Chem 292:18592–18607.

120. Padilla-Benavides T, Haokip DT, Yoon Y, Reyes-Gutierrez P, Rivera-Pérez JA, Imbalzano AN. 2020. CK2-Dependent Phosphorylation of the Brg1 Chromatin Remodeling Enzyme Occurs during Mitosis. Int J Mol Sci 21:1–21.

121. Giacinti C, Bagella L, Puri PL, Giordano A, Simone C. 2006. MyoD recruits the cdk9/cyclin T2 complex on myogenic-genes regulatory regions. J Cell Physiol 206:807–813.

122. Joliot V, Ait-Mohamed O, Battisti V, Pontis J, Philipot O, Robin P, Ito H, Ait-Si-Ali S. 2014. The SWI/SNF Subunit/Tumor Suppressor BAF47/INI1 Is Essential in Cell Cycle Arrest upon Skeletal Muscle Terminal Differentiation. PLoS One 9:1–11.

123. Serra C, Palacios D, Mozzetta C, Forcales S V., Morantte I, Ripani M, Jones DR, Du K, Jhala US, Simone C, Puri PL. 2007. Functional Interdependence at the Chromatin Level between the MKK6/p38 and IGF1/PI3K/AKT Pathways during Muscle Differentiation. Mol Cell 28:200–213.

124. Ohkawa Y, Imbalzano AN. 2006. Skeletal muscle specification by myogenin. EMBO J 25:490–501.

125. Mallappa C, Nasipak BT, Etheridge L, Androphy EJ, Jones SN, Sagerstro CG, Ohkawa Y, Imbalzano AN. 2010. Myogenic MicroRNA Expression Requires ATP-Dependent Chromatin Remodeling Enzyme Function. Mol Cell Biol 30:3176–3186.

126. de la Serna IL, Carlson KA, Hill DA, Guidi CJ, Stephenson RO, Sif S, Kingston RE, Imbalzano AN. 2000. Mammalian SWI-SNF Complexes Contribute to Activation of the hsp70 Gene. Mol Cell Biol 20:2839–2851.

127. Abràmoff MD, Magalhaes PJ, Ram SJ. 2004. Image Processing with ImageJ. Biophotonics Int 11:36–42.

128. Livak KJ, Schmittgen TD. 2001. Analysis of relative gene expression data using real-time quantitative PCR and the 2-ΔΔCT method. Methods 25:402–408.

129. Huang J, Liang X, Xuan Y, Geng C, Li Y, Lu H, Qu S, Mei X, Chen H, Yu T, Sun N, Rao J, Wang J, Zhang W, Chen Y, Liao S, Jiang H, Liu X, Yang Z, Mu F, Gao S. 2017. A reference human genome dataset of the BGISEQ-500 sequencer. Gigascience 6:1–9.

130. Kim D, Langmead B, Salzberg SL. 2015. HISAT: A fast spliced aligner with low memory requirements. Nat Methods 12:357–360.

131. Pertea M, Pertea GM, Antonescu CM, Chang TC, Mendell JT, Salzberg SL. 2015. StringTie enables improved reconstruction of a transcriptome from RNA-seq reads. Nat Biotechnol 33:290–295.

132. Trapnell C, Williams BA, Pertea G, Mortazavi A, Kwan G, Van Baren MJ, Salzberg SL, Wold BJ, Pachter L. 2010. Transcript assembly and quantification by RNA-Seq reveals unannotated transcripts and isoform switching during cell differentiation. Nat Biotechnol 28:511–515.

133. Langmead B, Salzberg SL. 2012. Fast gapped-read alignment with Bowtie 2. Nat Methods 9:357–359.

134. Li B, Dewey CN. 2011. RSEM: accurate transcript quantification from RNA-Seq data with or without a reference genome Bo. BMC Bioinformatics 12:1–16.

135. Love MI, Huber W, Anders S. 2014. Moderated estimation of fold change and dispersion for RNA-seq data with DESeq2. Genome Biol 15:1–21.

136. Audic S, Claverie JM. 1997. The significance of digital gene expression profiles. Genome Res 7:986–995.

