## Supplemental Figures for "The Bromodomains of the mammalian SWI/SNF (mSWI/SNF) ATPases Brahma (BRM) and Brahma Related Gene 1 (BRG1) promote chromatin interaction and are critical for skeletal muscle differentiation"

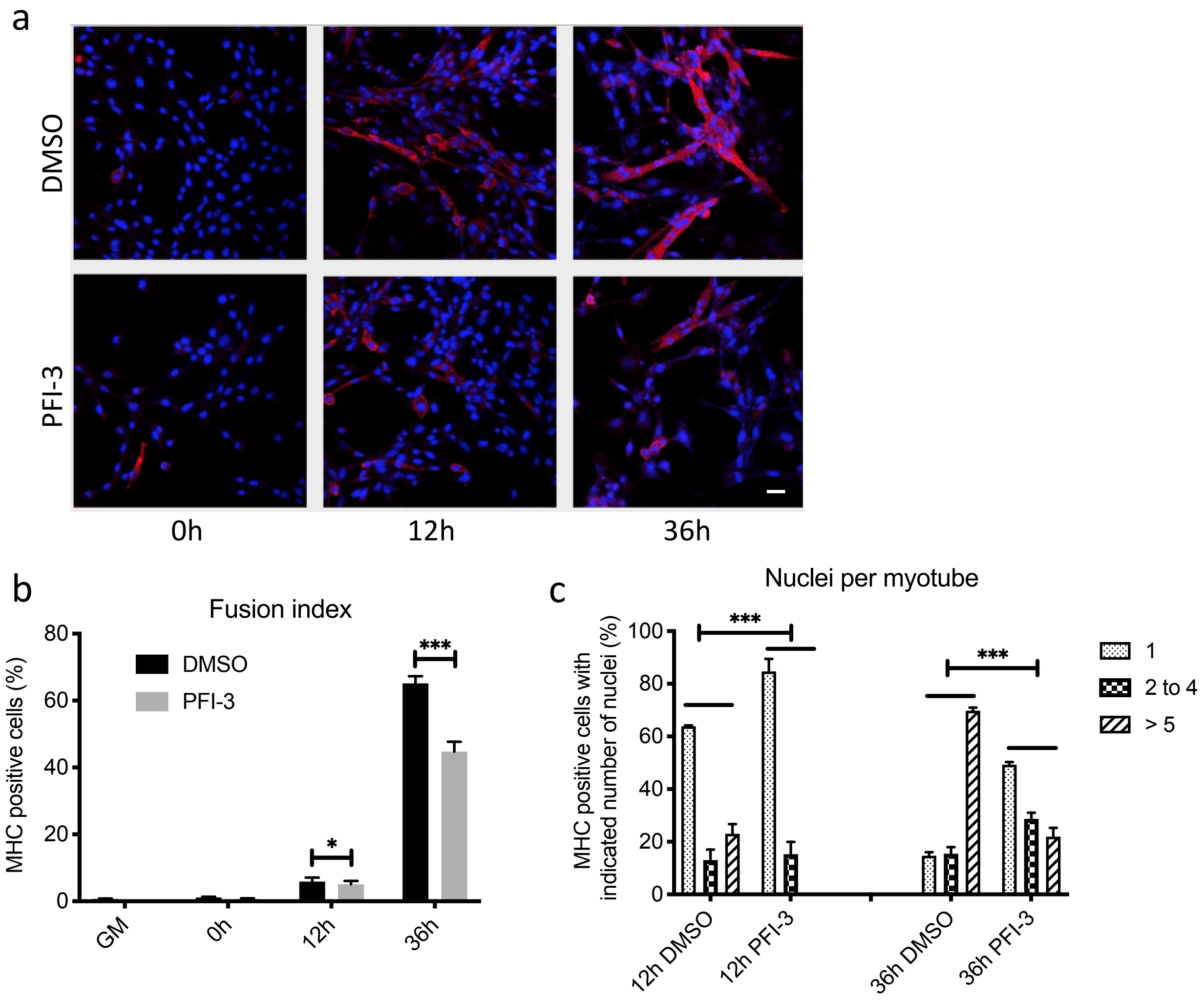


**Supp. Fig. 1:** (a) Confocal images for C57BL/6 mouse myoblasts treated with DMSO or PFI-3 and stained for myosin heavy chain (red) and nuclei (blue) at indicated timepoints, Scale 5µm. (b) Quantification of fusion index. (c) Differentiated myoblasts at the indicated timepoints were analyzed for number of nuclei per myotube. $*p<0.05$, $**p<0.01$ and $***p<0.005$

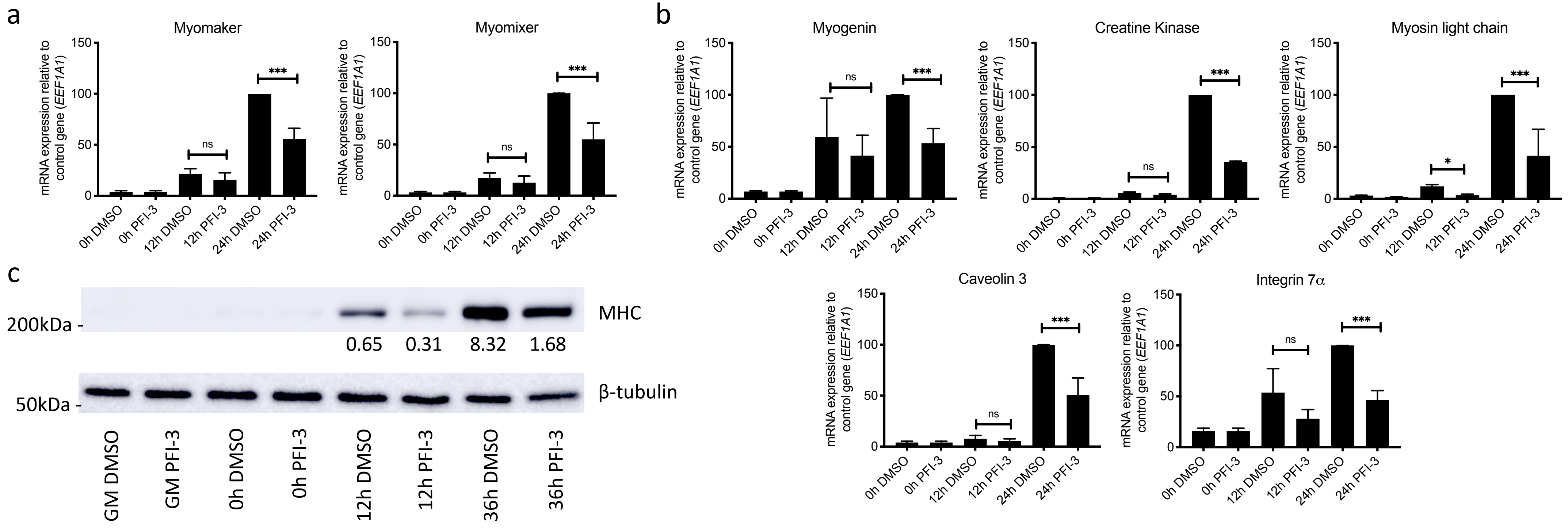


**Supp. Fig. 2:** (a) mRNA expression at the indicated timepoints for fusion regulator genes Myomaker and myomixer and (b) the myogenic genes Myogenin, muscle creatine kinase, myosin light chain 1, Caveolin 3 and Integrin 7α in C57BL/6 myoblasts treated with DMSO or PFI-3. Expression was normalized to a control gene (EEF1A1). 100% expression is defined as the timepoint at which maximal expression was observed. ns, not significant, $*p<0.05$, $**p<0.01$ and $***p<0.005$. (c) Representative western blot for MHC expression at the indicated times in C57BL/6 myoblasts treated with DMSO or PFI-3. The indicated numbers are the pixel counts normalized to β-tubulin expression calculated using ImageJ.

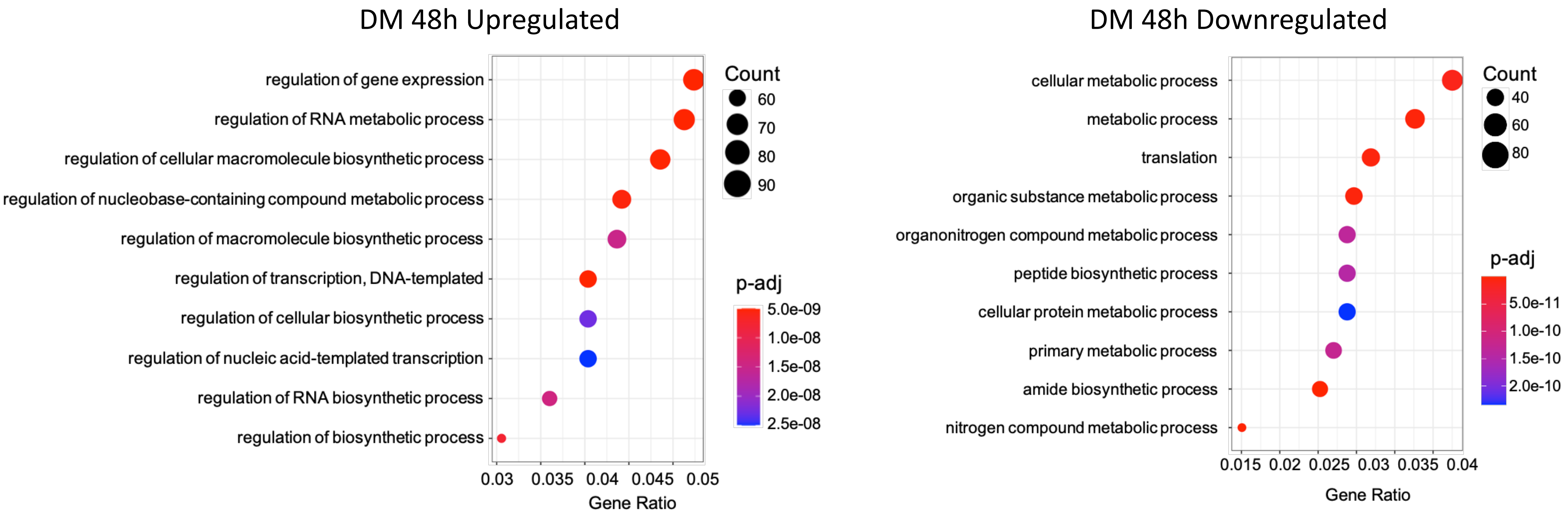


**Supp. Fig 3** GO analysis of BRG1-dependent genes as identified by Zhu et al (NAR, 2020) that not identified as PFI-3 dependent in our RNA-seq dataset.

a

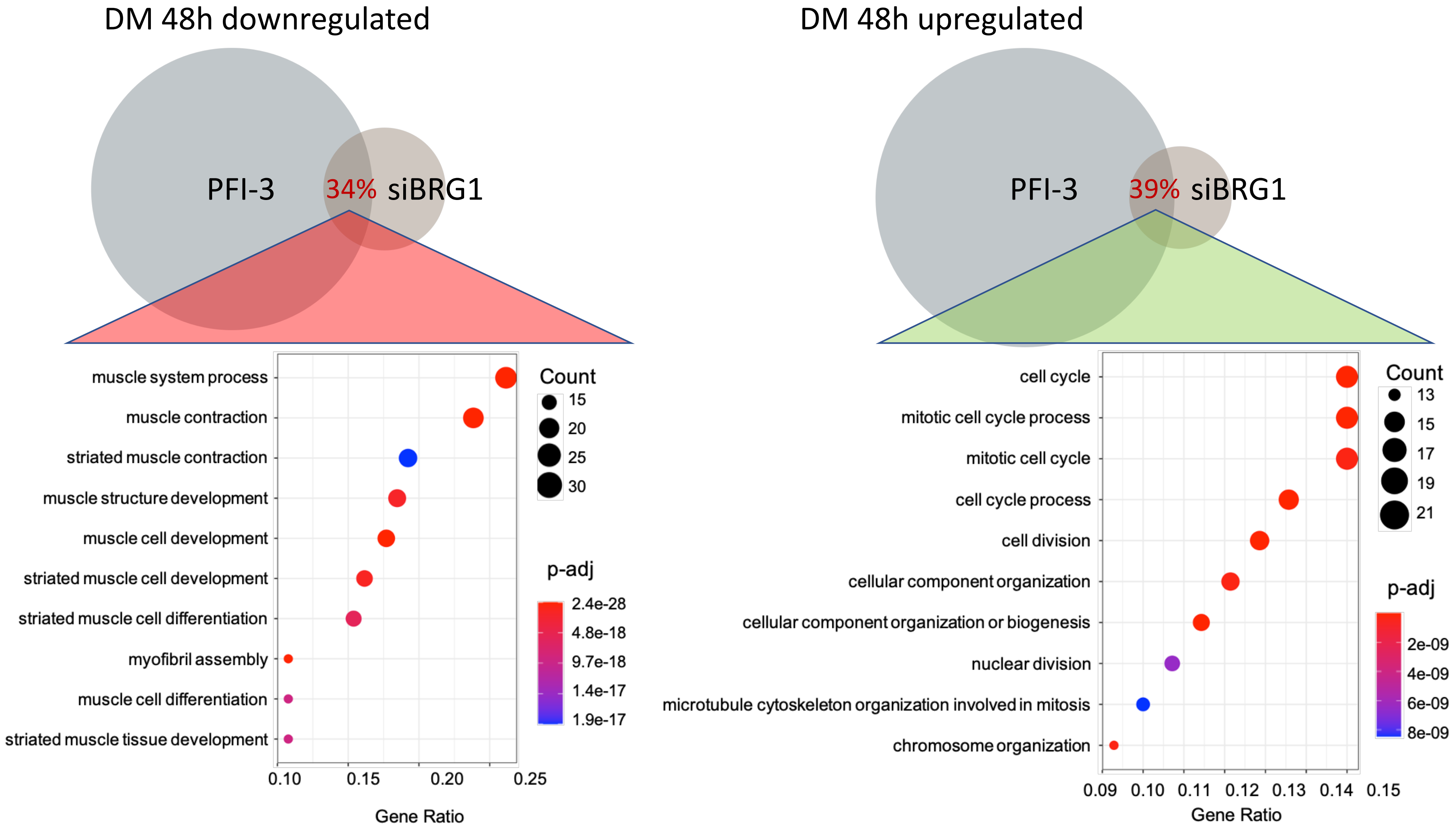


b

| Percentage overlap for DEGs between dataset from Albini et. al. and PFI-3 RNA-seq | | |
|---|---|---|
| Affected genes | Up | Down |
| Unique BRG1 targets | 39.0% | 33.6% |
| Unique BRM targets | 14.7% | 5.5% |
| BRG1/BRM co-regulated | 32.4% | 15.6% |

**Supp. Fig 4** (a) GO analysis of common target genes between the BRG1 knockdown dataset from Albini et. al. (EMBO Reports, 2015) and the PFI-3 treatment datasets shows upregulation of cell-cycle related genes and downregulation of muscle-related genes demonstrating the importance of BRG1 bromodomain function. (b) Comparative analysis showing the percentage of genes that are commonly regulated by BRG1 and/or BRM knockdown and PFI-3 treatment.
